# An open resource for transdiagnostic research in pediatric mental health and learning disorders

**DOI:** 10.1101/149369

**Authors:** Lindsay M. Alexander, Jasmine Escalera, Lei Ai, Charissa Andreotti, Karina Febre, Alexander Mangone, Natan Vega Potler, Nicolas Langer, Alexis Alexander, Meagan Kovacs, Shannon Litke, Bridget O’Hagan, Jennifer Andersen, Batya Bronstein, Anastasia Bui, Marijayne Bushey, Henry Butler, Victoria Castagna, Nicolas Camacho, Elisha Chan, Danielle Citera, Jon Clucas, Samantha Cohen, Sarah Dufek, Megan Eaves, Brian Fradera, Judith Gardner, Natalie Grant-Villegas, Gabriella Green, Camille Gregory, Emily Hart, Shana Harris, Megan Horton, Danielle Kahn, Katherine Kabotyanski, Bernard Karmel, Simon P. Kelly, Kayla Kleinman, Bonhwang Koo, Eliza Kramer, Elizabeth Lennon, Catherine Lord, Ginny Mantello, Amy Margolis, Kathleen R. Merikangas, Judith Milham, Giuseppe Minniti, Rebecca Neuhaus, Alexandra Nussbaum, Yael Osman, Lucas C. Parra, Ken R. Pugh, Amy Racanello, Anita Restrepo, Tian Saltzman, Batya Septimus, Russell Tobe, Rachel Waltz, Anna Williams, Anna Yeo, Francisco X. Castellanos, Arno Klein, Tomas Paus, Bennett L. Leventhal, R. Cameron Craddock, Harold S. Koplewicz, Michael P. Milham

## Abstract

Technological and methodological innovations are equipping researchers with unprecedented capabilities for detecting and characterizing pathologic processes in the developing human brain. As a result, ambitions to achieve clinically useful tools to assist in the diagnosis and management of mental health and learning disorders are gaining momentum. To this end, it is critical to accrue large-scale multimodal datasets that capture a broad range of commonly encountered clinical psychopathology. The Child Mind Institute has launched the Healthy Brain Network (HBN), an ongoing initiative focused on creating and sharing a biobank of data from 10,000 New York area participants (ages 5-21). The HBN Biobank houses data about psychiatric, behavioral, cognitive, and lifestyle phenotypes, as well as multimodal brain imaging (resting and naturalistic viewing fMRI, diffusion MRI, morphometric MRI), electroencephalography, eye-tracking, voice and video recordings, genetics, and actigraphy. Here, we present the rationale, design and implementation of HBN protocols. We describe the first data release (n = 664) and the potential of the biobank to advance related areas (e.g., biophysical modeling, voice analysis).

## 1. PURPOSE OF DATA COLLECTION

Psychiatric and learning disorders are among the most common and debilitating illnesses across the lifespan. Epidemiologic studies indicate that 75% of all diagnosable psychiatric disorders begin prior to age 24^1^. This underscores the need for increased focus on studies of the developing brain^2^. Beyond improving our understanding of the pathophysiology that underlies the emergence of psychiatric illness throughout development, such research has the potential to identify clinically useful markers of illness that can improve the early detection of pathology and guide interventions. Although the use of neuroimaging, neuropsychology, neurophysiology and genetics has made significant strides in revealing biological correlates for a broad array of illnesses, findings have been lacking in specificity^3^. Consequently, progress in finding clinically useful brain-based biomarkers has been disappointing^4,5^.

Given the slow pace in biomarker identification, investigators have been prompted to rethink research paradigms and practices. Most notably, the emphasis on mapping diagnostic labels from a clinically defined nosology (e.g., the Diagnostic and Statistical Manual of Mental Disorders (DSM) or the International Classification of Diseases) to varying biological indices has proven to be problematic, as it assumes consistent biological relationships with broad constellations of signs and symptoms^6,7^. Epidemiologists, psychopathologists, geneticists and neuroscientists are reconsidering the relevance of diagnostic boundaries due to the lack of specificity in related findings. Two psychiatric research approaches have emerged. First is the adoption of transdiagnostic models organized around behavioral and neurobiological dimensions that transcend existing diagnostic boundaries^3^. Second is the use of diagnostic subtyping to explain variation within diagnostic categories through the detection of behaviorally or biologically homogeneous subgroups^8–10^. These two strategies of parsing psychiatric illness are not mutually exclusive, and can inform each other.

Transdiagnostic and subtyping strategies call for changing the designs of future studies away from those typically applied to clinical neuroscience research^4^. First, we must move away from studying disorders in isolation from one another and from relying on *“extreme comparisons”* in which clinical samples are compared to healthy controls (often “super healthy” controls), rather than offering comparisons with individuals experiencing other clinical conditions. Unless this happens, the clinical relevance of published findings will remain limited, because they will provide little insight into real-world challenges of differentiating forms of psychopathology (i.e., a psychiatrist can easily differentiate an individual with schizophrenia from a healthy control but may find it much more challenging to determine whether psychosis or a mood disorder is the primary problem). Second, our science has been hindered by its reliance on small sample sizes that are vastly underpowered given the high dimensionality and usual small effect sizes of biological phenomena. This lesson is increasingly being incorporated in genetics, but it still applies to imaging or physiologically-based measures. Third, sample ascertainment can no longer be dependent on clinics, as the resulting samples bring with them a wide-range of multifaceted biases, including but not limited to symptom severity, sex distribution and problems related to access to care. As a result, there is a pressing need for community-based and epidemiologic samples^11^. Clearly, the time has come for changes in methods at all levels of psychiatric science.

In response to these challenges, and the scarcity of transdiagnostic datasets available for neuroscientific studies in children and adolescents, the Child Mind Institute has launched the Healthy Brain Network (HBN) initiative. As part of this initiative, the HBN is creating a Biobank from a community sample of 10,000 children and adolescents (ages 5-21) residing in the New York City area. The HBN Biobank includes behavioral and cognitive phenotyping, as well as multimodal brain imaging, electroencephalography (EEG), eye tracking, genetics, digital voice and video samples, and actigraphy. The HBN Biobank has an extensive phenotyping protocol that includes comprehensive psychiatric and learning assessments, as well as instruments probing a range of familial, environmental and lifestyle variables (e.g., physical activity, nutrition). Consistent with the model established by the NKI-Rockland Sample^12^, all data obtained are being shared on a pre-publication basis throughout the six-year course of the data acquisition phases for the project. Taken together, access to such a range of data will ensure that the HBN Biobank will allow for scholars to address rich and clinically relevant questions.

What follows is an overview of the project plan and protocol details for the HBN Biobank; we also describe strategies and tests developed as part of the process of ensuring that the HBN initiative can be scaled up to meet its high throughput goals. Finally, we provide descriptions and quality assurance characteristics for the initial major data release (n=664).

## 2. METHODS

### 2.1 Recruitment Strategy

A primary goal for the HBN is to generate a dataset that captures the broad range of heterogeneity and impairment that exists in developmental psychopathology. Accordingly, we adopted a community-referred recruitment model. We use advertisements to encourage participation of families who have concerns about psychiatric symptoms in their child. The “announcements” are distributed to community members, educators and local care providers, as well as directly to parents via email lists and events. The advertisements highlight the potential value of participation for children who may require school-based accommodations. In particular, the comprehensive diagnostic evaluation reports provided by HBN include clinical impressions and actionable treatment recommendations; when appropriate, the reports can be used to acquire an Individualized Education Program (IEP) - a prerequisite for obtaining school accommodations, services, and specialized classroom placements. Upon completion of the study, we offer participants referral information and up to to three in-person feedback sessions. Modest monetary compensation for their time and expenses incurred are also provided.

It is important to note that our recruitment strategy was developed to achieve the major goals of the HBN after considering the alternative of a fully representative epidemiologic design. The primary HBN goal is to generate a large-scale, transdiagnostic sample for biomarker discovery and for investigations of the neural substrates associated with commonly occurring illness phenotypes. While HBN ascertainment is not clinic-based, per se, the strategy of recruiting on the basis of perceived clinical concern dictates that the HBN sample will include a high proportion of individuals affected by psychiatric illness. Despite the lack of rigorous epidemiologic ascertainment, the intended scale of data collection and the inclusion of inherently diverse communities across NYC may approximate representativeness for the sample. The scale of the sample should also allow investigators to study selected sub-cohorts of interest for targeted study (e.g., comparing individuals with ADHD residing in Midtown Manhattan vs. those residing in Staten Island). Finally, depending on the ability to secure financial support, the fourth phase of HBN will switch strategies to make the final 1500 participants a representative epidemiologic cohort.

### 2.2 Participant Procedures

#### 2.2.1 Screening

To determine eligibility and ensure safety, potential participants or their legal guardians (if they are under age 18) complete a prescreening phone interview with an intake coordinator. This screening interview obtains information regarding a potential participant’s psychiatric and medical history. With few exceptions, the presence of psychiatric, medical, or neurological illness does not exclude participation. Primary causes for exclusion center on the presence of acute safety concerns (e.g., danger to self or others), cognitive or behavioral impairments that could interfere with participation (e.g., being nonverbal, IQ less than 66) or medical concerns that are expected to confound brain-related findings (see Table 1). All individuals meeting inclusion criteria, without any reasons for exclusion, are invited to participate in the study.

**Table 1.** Participant Inclusion and Exclusion Criteria.

| Inclusion Criteria |  |
| --- | --- |
| 1. | Male or female ages 5 - 21 years. |
| 2. | Adults must have capacity to understand the study and provide informed consent.<br>a. Children ages 5-17 must have the capacity to provide assent ( must speak in simple, but full (3+ word) sentences at the Kindergarten level) and parent/guardian must have the capacity to sign informed consent. |
| 3. | Participants must be fluent in English. Children who are fluent in English but have parents who speak Spanish can be enrolled upon availability of Spanish-speaking personnel. |
| Exclusion Criteria |  |
| 1. | Serious neurological (specific or focal) disorders preventing full participation in the protocol. Some children with moderate to severe impairment in cognitive (i.e., IQ below 66) and/or general function will be eligible for a short, research-based protocol. Parents will be informed that this battery will not allow for a full, comprehensive feedback report. A feedback session and abbreviated report will be provided.<br>a. Examples include: non-verbal and/or low functioning autism, placement in classroom environment lower than 12:1:1, chronic epilepsy. |
| 2. | Childhood metabolic disorders (i.e., disorders of lipid, carbohydrate, metal, amino acid, etc.). |
| 3. | Positive HIV status. |
| 4. | Renal or liver failure.<br>a. Must be current, acute failure. Past liver or kidney disease that was treated to remission confers eligibility. |
| 5. | Current or past diagnosis of encephalitis. |
| 6. | Known neurodegenerative disorder (e.g. Huntington’s Disease, ALS, MS, Cerebral Palsy). |
| 7. | Hearing or visual impairment that prevents participation in study-related tasks.<br>a. Child can participate if vision or hearing is corrected with devices. |
| 8. | Current history (within the past 6 months) of Schizophrenia, Schizoaffective Disorder, or Bipolar Disorder.<br>a. The absence of a formal diagnosis confers eligibility. |
| 9. | Manic or psychotic episode within the past 6 months without current, ongoing treatment. |
| 10. | New onset (within the last 3 months) of suicidality or homicidality for which there is no current, ongoing treatment.<br>a. This can be ideation or a plan. It must be believable, recurrent, and bona fide. |
| 11. | History of lifetime substance dependence requiring chemical replacement therapy. |
| 12. | History of substance dependence within the last year except nicotine and marijuana. |

#### 2.2.2 Medication

Participants taking stimulant medication are asked to discontinue their medication during the days of participation, as stimulants are known to have an effect on cognitive and behavioral testing, as well as functional brain mapping. Participants who choose not to discontinue medication, or whose physicians require that medication not be interrupted, are still enrolled. Medication taken on the day of participation is recorded.

#### 2.2.3 IRB Approval

The study was approved by the Chesapeake Institutional Review Board (https://www.chesapeakeirb.com/). Prior to conducting the research, written informed consent is obtained from participants ages 18 or older. For participants younger than 18, written consent is obtained from their legal guardians and written assent obtained from the participant.

### 2.3 Project Plan

The HBN has a four-phase project plan (see Table 2). The goals for each of the phases are as follows:

**Table 2.** Healthy Brain Network Project Plan.

| **Project Plan Goals** |  |
| --- | --- |
| **Phase I: Implementation and Testing (N=500)** |  |
| Establish project workflows<br>Mental health diagnostic evaluation<br>Phenotypic assessment<br>EEG<br>MRI scanning<br>Establish prototypes for Diagnostic Research Center (HBN-Staten Island) | Test utility of mobile Diagnostic Research Center<br>Test utility of mobile MRI platform<br>Establish recruitment sources and community partners |
| **Phase II: Revision and Hardening (N=500)** |  |
| Augment learning and language evaluation protocols<br>Increase breadth of phenotyping and overlap with other initiatives<br>Family history<br>Impairment<br>Prenatal assessment<br>Parental distress<br>Sleep<br>Stress/trauma<br>Substance use<br>Identify and troubleshoot data quality issues | Introduce voice assessment protocols<br>Introduce saliva collection for genetics<br>Introduce natural viewing fMRI<br>Introduce home-based longitudinal follow-up (HBN Quarterly Mental Health Report)<br>Optimize staffing models and workflow efficiencies<br>Test and harden KSADS-COMP<br>Test and harden E-SWAN<br>Test reproducibility of prototype Diagnostic Research Center (HBN-Manhattan) |
| **Phase III: Scale-up (N=7,500)** |  |
| Increase to three full-scale Diagnostic Research Centers<br>Transition to multi-site stationary MRI scanner model | Implement infrastructure for epidemiologic sampling<br>Introduce home-based phlebotomy collection model<br>Introduce actigraphy |
| **Phase IV: Targeted Recruitment (N=1,500)** |  |
| Epidemiologic sampling |  |

*<u>Phase I: Implementation and Testing (Participants 1-500; completed).</u>* The overarching goal of the initial phase was to establish a prototype HBN Diagnostic Research Center, located in Staten Island, New York (one of the five boroughs of New York City). Prototype development was intended to establish all project workflows and strategies/procedures for recruitment, diagnostic evaluations, phenotypic assessments, and a referral network (i.e., health care providers to whom participants can be referred if clinical significant concerns are detected). The initial protocol included diagnostic evaluations, phenotypic assessments, EEG and magnetic resonance imaging (MRI). During the initial phase, we also evaluated the feasibility and benefits of using a mobile MRI scanner, as well as a mobile Diagnostic Research Center.

*<u>Phase II: Revision and Hardening (Participants 501-1000; completed).</u>* A key challenge for almost any large-scale study is balancing the desire to maintain stable protocols and assessments across the entirety of a sample with the desire to integrate new measures and make changes based on learning from experiences and scientific advances along the way. Phase II of the Healthy Brain Network had two primary goals: 1) the addition and/or deletion of protocols established during Phase I, based on lessons learned and new developments; and, 2) hardening the revised protocols to ensure that they are as optimal and robust as possible, while also reflecting the current state of the art in science and practice.

*<u>Phase III: Scale-up (Participants 1001-8500; in process).</u>* Building on the experience and lessons learned from Phases I and II of the project, the Healthy Brain Network has started Phase III, with the goal of enrolling 7,500 participants in our established protocol. This goal necessitates increased capacity for both recruitment and enrollment. As such, Phase III includes additional Diagnostic Research Centers and MRI scan sites in the New York City region; sites are being chosen to increase the diversity of populations that can be reached.

*<u>Phase IV: Targeted Recruitment (Participants 8501-10000).</u>* The final phase of the Healthy Brain Network will incorporate epidemiologic sampling to recruit an additional targeted representative sample of 1,500 participants.

### 2.4 Experimental Design

The HBN protocol spans four sessions, each approximately three hours in duration (see Table 3). A list of all measures collected during the four-session evaluation can be found in Table 4. The assessment includes:

**Table 3.** HBN Visit Schedule, Manhattan and Staten Island Offices.

**Healthy Brain Network Visit Schedule**
| Visit One |  |  |
| --- | --- | --- |
| Time (min) | Child Activity | Parent Activity |
| 30 |  | Introduction, consent |
| 15 | Assent | Enrollment, MRI screening |
| 75 | WISC/WASI/KBIT | Pre-Interview I (clinical portion) |
| 30 | Child questionnaires |  |
| 15 | Mock scanner |  |

| Visit Two |  |  |
| --- | --- | --- |
| Time (min) | Child Activity | Parent Activity |
| 105 | MRI scan |  |

| Visit Three |  |  |
| --- | --- | --- |
| Time (min) | Child Activity | Parent Activity |
| 45-69 | WIAT | Pre-Interview II: RA portion |
| 20-30 | CELF-5 and TOWRE | Parent questionnaires |
| 30 | NIH Toolbox |  |
| 60 | Child questionnaires |  |
| 30 | Fitness/vitals |  |

| Visit Four |  |  |
| --- | --- | --- |
| Time (min) | Child Activity | Parent Activity |
| 75-90 | EEG | KSADS |
| 40-45 | KSADS |  |
| 30 | Quotient |  |
| 30 | CTOPP and GFTA |  |

**Table 4.** Complete HBN Protocol.

| General Information | Behavioral Measures |
| --- | --- |
| Demographics<br>CMI Symptom Checker<br>Edinburgh Handedness Inventory<br>Intake Interview<br>Physical Activity Questionnaire for Older Children (PAQ-C) (8-14)<br>Physical Activity Questionnaire for Adolescents (PAQ-A) (14-19)<br>Barratt Simplified Measure of Social Status<br>Financial Support Questionnaire<br>Medical History Questionnaire – Family<br>Pregnancy and Birth Questionnaire | Child Behavior Checklist (CBCL) (5-17)<br>Youth Self Report (YSR) (11-18)<br>Adult Self Report (ASR) (18+)<br>Screen for Child Anxiety Related Disorders (SCARED) – Parent Report & Self Report (8-18)<br>State Trait Anxiety Inventory (STAI) (18+) – Self Report<br>Mood & Feelings Questionnaire (MFQ) – Parent Report & Self Report (8+)<br>Affective Reactivity Index – (ARI-S) Self Report<br>Columbia Suicide Severity Rating Scale (C-SSRS) – Self Report (7+)<br>Extended Strengths and Weaknesses Assessment of Normal Behavior (E-SWAN) (5-17)<br>Strengths and Weaknesses of ADHD Symptoms and Normal Behavior Scale (SWAN) (6+)<br>Conners ADHD Rating Scales Self Report Short Form (Conners) (8+)<br>Repetative Behavior Scale (RBS) (5-21)<br>Autism Spectrum Screening Questionnaire (ASSQ) (5+)<br>Social Communication Questionnaire (SCQ) (5+)<br>Social Responsiveness Scale-2 (SRS-2) (5+)<br>Strengths and Difficulties Questionnaire (5+)<br>The Columbia Impairment Scale (CIS) Parent and self report (5+)<br>Social Aptitudes Scale (SAS) (5+)<br>WHO Disability Assessment Schedule (WHODAS) Parent and Self-Report (5+)<br>Food Frequency Questionnaire (FFQ) (5-17)<br>Inventory of Callous-Unemotional Traits - Parent Report (5+) |
| Physical Measures | Family Structure, Stress and Trauma |
| FITNESSGRAM (Pushups, Curl-ups, Trunk-Lift, Sit and Reach, Grip Strength)<br>Cardiovascular Fitness Test<br>Vitals (Heart Rate, Blood pressure)<br>Measurements (Height/weight, Waist circumference, Bio-impedance)<br>Blood Draw (Endocrine, Immunologic, and Metabolic profiling; Genetics)<br>Buccal Swabs (Genetics)<br>Urine Sample (Toxicology screen, Pregnancy test: 11+)<br>Ishihara Color Vision Test<br>Electroencephalography (EEG)/Eye Tracking<br>Magnetic Resonance Imaging (MRI)<br>Peterson Puberty Scale (6-17)<br>Sleep Disturbance Scale for Children (SDSC) (6-15) | Family History-Research Diagnostic Criteria (FH-RDC)<br>Parental Stress Index IV (PSI-IV)<br>Alabama Parenting Questionnaire – Self Report (APQ) (6-18)<br>Alabama Parenting Questionnaire – Parent Report (APQ) (6-18)<br>Children's Perception of Interparental Conflict (CPIC) (8-18)<br>Distress Tolerance Index – Parental Self Report<br>Children's Coping Strategies Checklist – Revised (CCSC) (8-18)<br>UCLA Trauma Reactivity Sale for DSM-V (UCLA) (5-18)<br>Negative Life Events Scale (NLES) – Self Report (8-18)<br>Negative Life Events Scale Parent Report (8-18)<br>Adverse Childhood Experiences Scale (ACES) (18+) |
| Cognition and Language Tasks | Substance Use and Addiction Measures |
| NIH Toolbox Tasks: Flanker, Card Sort and Processing Speed<br>Temporal Discounting Task<br>Quotient ADHD System<br>Rapid Automatic Naming & Rapid Alternating Stimulus Test (RAN/RAS) (5)<br>Wechsler Intelligence Scale for Children-V (WISC-V) (6-17)<br>Wechsler Adult Intelligence Scale-IV (WAIS-IV): (17+)<br>Wechsler Abbreviated Scale of Intelligence-II (WASI): (17+)<br>Wechsler Individual Achievement Test – III (WIAT)<br>Differential Ability Scales – II (DAS) (5 or IQ below 70)<br>Clinical Evaluation of Language Fundamentals – 5th Edition (CELF-5)<br>Goldman Fristoe Test of Articulation – II (GFTA)<br>Comprehensive Test of Phonological Processing – II (CTOPP)<br>Test of Word Reading Efficiency (TOWRE) (6+)<br>Expressive Vocabulary Test (EVT) (when indicated)<br>Peabody Picture Vocabulary Test (PPVT) (when indicated) | Fagerstrom Test for Nicotine Dependence (FTND) (18+)<br>Alcohol Use Disorders Identification Test (AUDIT) (11+)<br>Modified Fagerstrom Tolerance Questionnaire- Adolescents (FTQA) (13-17)<br>European School Survey Project on Alcohol & Other Drugs (ESPAD) (10+)<br>Internet Addiction Test (IAT)<br>Parent-Child Internet Addiction Test (PCIAT)<br>Yale Food Addiction Scale (YFAS) and YFAS-Child |
| Diagnostic Assessments | Longitudinal Follow Up Measures |
| Kiddie Schedule for Affective Disorders and Schizophrenia (K-SADS)<br>Child and Adolescent Psychiatric Assessment Schedule (Cha-PAS) (when indicated)<br>Vineland Adaptive Behavior Scale – Parent/Caregiver Rating Form (when indicated)<br>Yale Global Tic Severity Scale (YGTSS) (when indicated, 6+)<br>Yale-Brown Obsessive Compulsive Scale (Y-BOCS) (when indicated, 18+)<br>Children's Yale-Brown Obsessive Compulsive Scale (when indicated, 6-18) | Youth Services Survey (YSS) & Services Assessment for Children and Adolescents (SACA)<br>Follow Up: CBCL<br>Follow Up: Columbia Impairment Scale Parent and Self Report<br>Follow Up: WHODAS Parent and Self Report |

#### 2.4.1 Clinician-Administered Assessments

The clinical staff consists of a combination of psychologists and social workers, with psychopharmacological consultation support provided by psychiatrists. All the tests in this section are administered by, or directly under the supervision of, licensed clinicians. Participant responses are first scored by the administering clinician. To enhance validity, the entire set of responses is again scored by a trained research assistant. Finally, all test scores from clinical interviews are double-entered into the database by two (different) trained research assistants.

*<u>Semi-Structured Diagnostic Interview.</u>* All participants are administered a computerized web-based version of the Schedule for Affective Disorders and Schizophrenia - Children’s version (KSADS)^13^. The KSADS-COMP is a semi-structured DSM-5-based psychiatric interview used to derive clinical diagnoses; administration in the HBN is performed by a licensed clinician. The KSADS-COMP includes a clinician-conducted parent interview and child interview, which result in automated diagnoses. Following completion of the interviews and review of all materials collected during study participation, clinically synthesized diagnoses (i.e., consensus DSM-5 diagnoses) are generated by the clinical team. The HBN data include the KSADS-COMP interview data along with the algorithm-generated diagnoses, as well as consensus clinical diagnoses, for each participant.

*<u>Additional Diagnostic Assessments.</u>* For a subset of psychiatric disorders, specific follow-up assessments are completed, as indicated for additional clinical characterization beyond the KSADS (e.g., Autism Diagnostic Observation Schedule [ADOS]^14^ for suspected autism, Clinical Evaluation of Language Fundamentals [CELF]^15^ for suspected language disorder) (See Table 5). These targeted supplemental diagnostic assessments are not administered to individuals without a suspicion of the presence of clinically significant illness in the corresponding domain.

**Table 5.** HBN Diagnostic Specific Assessments.

| **Diagnosis** | **Assessment** | **Description** |
| --- | --- | --- |
| **Autism Spectrum Disorder** | Autism Diagnostic Interview – Revised (ADI-R) | A reliable and valid standardized diagnostic interview developed to aid practitioners in gathering a complete developmental history and current functioning level for an individual being evaluated for ASD. Administered to participants with a referral for autism-specific evaluation. |
|  | Autism Diagnostic Observation Schedule, 2<sup>nd</sup> edition (ADOS-2) | A standardized, semi-structured play-based assessment in which tasks are presented in a standardized manner to elicit and/or highlight the presence or absence of specific behaviors relevant to making an ASD diagnosis. Administered to participants with a referral for autism-specific evaluation. |
| **Intellectual Disability** | Vineland Adaptive Behavior Scale – Parent/Caregiver Rating Form | A measure of adaptive behavior from birth to adulthood; forms an aid in diagnosing and classifying intellectual and developmental disabilities. Administered to parents of participants with developmental or intellectual disorders. |
|  | Child and Adolescent Psychiatric Assessment Schedule (ChA-PAS) | A semi-structured clinical interview linked to a clinical glossary that guides the ratings. The ChA-PAS has, however, been extended to include ADHD and Behavioral Disorders, as well as axis I psychiatric disorders. It also includes a screen for autistic spectrum disorders. Administered to parents of participants with developmental or intellectual disorders. |
| **Speech/Language Disorder** | Clinical Evaluation of Language Fundamentals – Fifth Edition (CELF-5) | An individually administered assessment tool made up of 18 subtests organized into four levels of testing that address language content, structure, and use. Administered to children with a referral for an extended language evaluation. |
|  | Test of Language Competence – Expanded Edition (TLC-E) Level 1 | An individually administered, norm-referenced oral language measure which evaluates for delays in the emergence of linguistic competence and in the use of semantic, syntactic, and pragmatic-strategies. An emphasis is placed on assessing within the contextual and situational demands of conversation in addition to basic semantic and syntactic abilities. Administered to children with a referral for an extended language evaluation. |
|  | Expressive Vocabulary Test, Second Edition (EVT-2) | An individually administered, norm-referenced instrument that assesses expressive vocabulary and word retrieval for children and adults. Administered to children with a referral for an extended language evaluation. |
|  | Peabody Picture Vocabulary Test, Fourth Edition (PPVT-4) | A norm-referenced, wide-range instrument for measuring the receptive (hearing) vocabulary of children and adults. For each item, the examiner says a word, and the examinee responds by selecting the picture that best illustrates that word's meaning. Administered to children with a referral for an extended language evaluation. |
|  | Goldman-Fristoe Test of Articulation - III (GFTA-3) | Provides information about a child's articulation ability by sampling both spontaneous and imitative sound production. Use this test to measure articulation of consonant sounds, determine types of misarticulation, and compare individual performance to national, gender-differentiated norms. Administered to children with a referral for an extended language evaluation. |
| **Obsessive Compulsive Disorder (OCD)** | Yale-Brown Obsessive Compulsive Scale (Y-BOCS) | A semi-structured clinician-rated instrument that assesses the severity of OCD symptoms. Administered as part of KSADS interview, if indicated. |
| **Tic Disorder** | Yale Global Tic Severity Scale (Y-GTSS) | A semi-structured clinician-rated instrument that assesses the nature of motor and phonic tics. Administered as part of KSADS interview, if indicated. |

*<u>Intelligence and Learning.</u>* Participants ages 6-17 complete the Wechsler Intelligence Scale for Children (WISC-V)^16^. Participants age 5, and those believed to have an IQ below 70, complete the Kaufman Brief Intelligence Test (KBIT)^17^. Participants ages 18 and older complete the Wechsler Adult Intelligence Scale (WAIS-IV)^18^. All participants ages 6 and older complete the Wechsler Individual Achievement Test (WIAT III)^19^.

*<u>Language.</u>* Trained research assistants and clinicians administer language screening tests as indicated, including the Clinical Evaluation of Language Fundamentals (CELF-5) Screener, the Goldman Fristoe Test of Articulation (GFTA) ‘Sounds and Words’ subtest^20^, the Comprehensive Test of Phonological Processing, Second Edition (CTOPP-2)^21^, and the Test of Word Reading Efficiency, Second Edition (TOWRE-2) Evaluation of Language Fundamentals [CELF]^21^, and the Test of Word Reading^22^. In addition, participants who fail the CELF-5 Screener and/or perform poorly on GFTA subtests are offered additional language evaluations performed by a licensed speech and language pathologist. This assessment includes the full CELF-5 assessment^23^, Expressive Vocabulary Test (EVT)^24^, the Peabody Picture Vocabulary Test (PPVT)^25^, the CELF-5 Metalinguistics^23^, and additional subtests of the GFTA.

#### 2.4.2 Self-Administered Assessments

Participant report and parent measures are acquired via the online patient portal of the NextGen electronic medical record system. Direct electronic entry of responses by participants minimizes the burden on research staff and removes the potential for errors that arise when questionnaires are administered using pen and paper, and then manually entered into a database. Structured questionnaires assess behavior, family structure, stress and trauma, as well as substance use and addiction (see Table 4). Each participant completes a set of questionnaires specific for his/her age and according to the protocol version at time of participation. See Figure 1 for a timeline of changes to the HBN assessment protocol over the first two years of the project.

**Figure 1.**
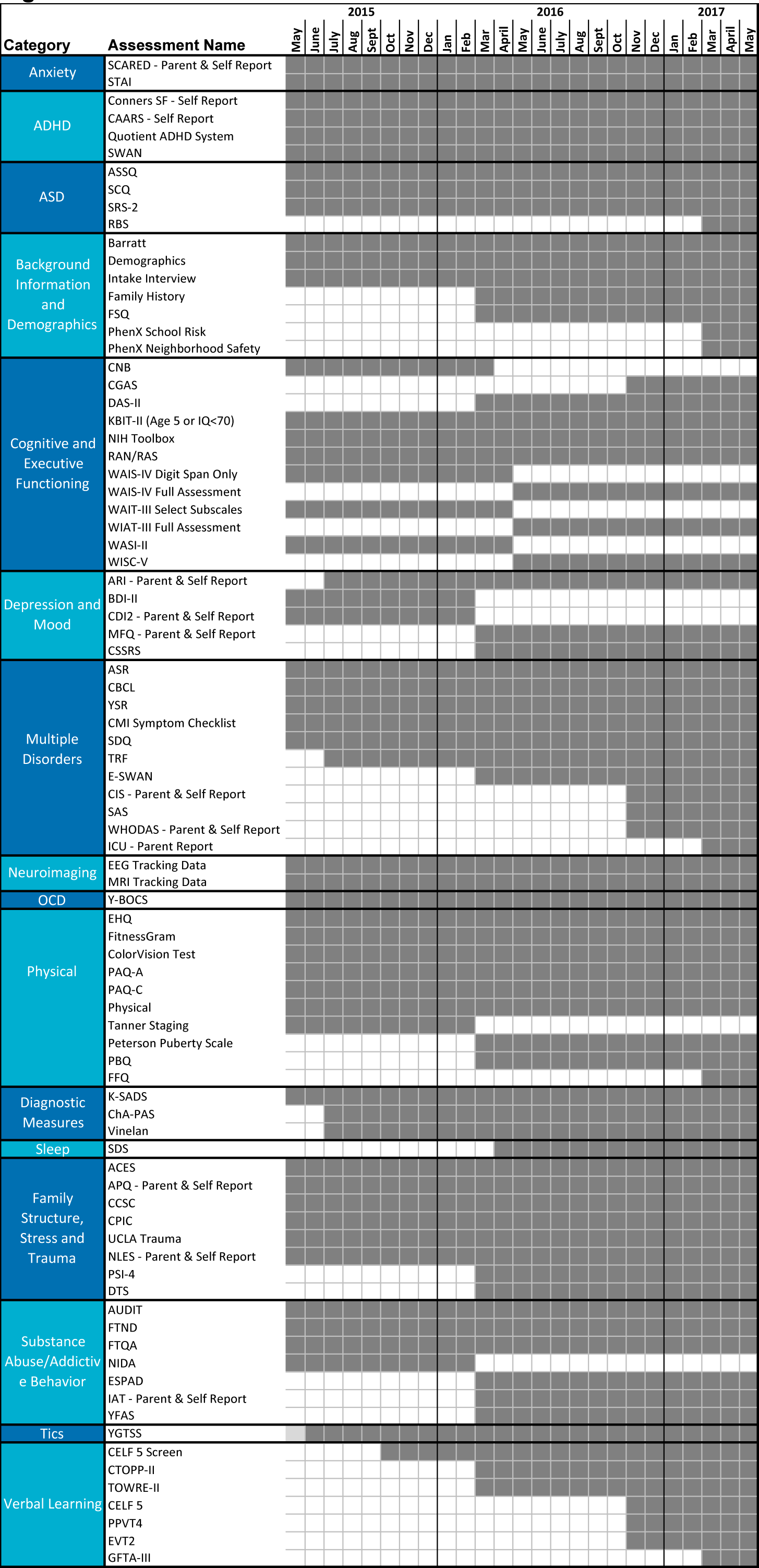
*HBN Protocol Timeline.* Here we depict the month in which each assessment was added (and in some cases removed). Dark gray boxes indicate inclusion of the assessment in the protocol for a given month, while white boxes indicate the measure was not included.

In the case of teacher reports, paper forms are used to collect data (e.g., Teacher Report Forms^26^) due to varying levels of receptiveness for electronic forms. All data collected on paper are double-entered by trained research assistants.

#### 2.4.3 Computerized Testing

Given the emphasis on clinically and educationally relevant assessments, limited time was available for additional computerized testing. To facilitate overlaps with cognitive phenotyping in other efforts, a subset of the NIH Toolbox has been included, consisting of: Flanker Task (Executive Function/Inhibitory Control and Attention), Card Sort (Executive Function/Dimensional Change), and Pattern Comparison (Processing Speed)^27^. In June 2017, an additional 1-minute task measuring temporal discounting was added to the HBN protocol^28^.

#### 2.4.4 Fitness Testing

Basic physical measurements (e.g., height, weight, and waist circumference) and cardiovascular measures (e.g., blood pressure and heart rate) are collected by trained research assistants. Cardiovascular fitness is assessed using a modified version of the FitnessGram test battery. FitnessGram^29^ is a widely used health-related physical fitness assessment that measures five different parameters, including aerobic capacity, muscular strength, muscular endurance, flexibility, and body composition. A treadmill test is used to measure maximal oxygen consumption for the purposes of estimating VO_2_max. Bioelectric impedance measures, used for the calculation of various indices of body composition (e.g., body mass index,percent body fat, percent water weight), are taken using the RJL Systems Quantum III BIA system.

#### 2.4.5 Electroencephalography (EEG) and Eye Tracking

For each participant, EEG and eye-tracking data are obtained during a battery that was previously assembled to examine attention, perception, inhibitory control, and decision-making^30^. See Table 6 for the specific paradigms and brief descriptions of each.

**Table 6.** Description of EEG Paradigms.

| Task | Degree of stimulation | Description | Duration | Reference* |
| --- | --- | --- | --- | --- |
| **Active (task-dependent paradigms)** |  |  |  |  |
| Sequence Learning Paradigm | Moderate | Participants are asked to observe and memorize of a sequence of either five or ten stimuli, depending on age. The sequence is repeated across five trials.<br>The purpose of the paradigm is to track the progress of gradual memory formation. EEG and/or ERP signatures of basic stimulus processing and memory encoding can be examined with respect to behavioral indices of learning performance on a block-by-block or trial-by-trial basis. | 5 mins | Steinemann, N. A., Moissello, C., Ghilardi, M. F., & Kelly, S. P. (2016). Tracking neural correlates of successful learning over repeated sequence observations. *NeuroImage*, 137, 152-164. |
| Visual Perception/Decision-making Paradigm | Moderate | Participants continuously monitor two overlaid, flickering grating patterns on a screen, indicating with a button press when they detect a contrast difference between the two.<br>This task furnishes behavioral metrics (reaction time, accuracy) for simple decisions and electrophysiological signatures of evidence encoding, accumulation, and motor preparation. | 9 mins<br>(3 runs of 3 minutes each) | O'Connell, R. G., Dockree, P. M. & Kelly, S. P. A supramodal accumulation-to-bound signal that determines perceptual decisions in humans. *Nature neuroscience* 15, 1729–1735 (2012). |
| WISC-IV Symbol Search Paradigm | Complex | For each trial in the symbol search paradigm, participants are shown rows of two target symbols and five symbols, and asked to indicate whether or not one of the target symbols appears in one of the five subsequent symbols.<br>The paradigm is a computerized version of a clinical pediatric assessment intended to measure processing speed capacity. Eye tracking is used to gather information about how long participants look at each symbol and their strategy for completing the task. | 2 mins | Wechsler, D. The Wechsler intelligence scale for children. 4th edn (Pearson, 2004). |
| **Passive (task-independent paradigms)** |  |  |  |  |
| Resting-State | None | Participants view a fixation cross on the center of the computer screen. Throughout the paradigm, participants are instructed to open or close their eyes at various points.<br>The paradigm is intended to measure endogenous brain activity during rest. | 5 mins | Fox, M. D. & Greicius, M. Clinical applications of resting state functional connectivity. *Frontiers in systems neuroscience* 4, 19 (2010). |
| Inhibition/Excitation Paradigm | Minimal | The stimulus used for this paradigm consists of four small flickering discs embedded in a static grating background. The discs generate strong steady-state responses that vary with contrast of the flickering stimuli.<br>The paradigm is intended to measure excitatory (SSVEP) and inhibitory (surround suppression) neurophysiological activity. | 3.5 mins | Vanegas, M. I., Blangero, A., & Kelly, S. P. (2015). Electrophysiological indices of surround suppression in humans. *Journal of neurophysiology*, 113(4), 1100-1109. |
| Naturalistic Stimuli Paradigm | Complex | Participants view a montage of short video clips taken from age-appropriate, mainstream television and movies.<br>Stimuli include the following:<br>*Despicable Me* (Clip from feature-length film; 3 mins)<br>*Diary of a Wimpy Kid* (Trailer for feature-length film; 2 mins)<br>"Fun with Fractals" (Educational video clip; 4.5 mins)<br>*The Present* (Short film; 4.5 mins)<br>The purpose of this paradigm is to measure neurophysiological activity during higher-level audio-visual stimulation. | 14 mins | Hasson, U., et al. Intersubject synchronization of cortical activity during natural vision. *Science* 303, 1634–1640 (2004).; Hasson, U., et al. Reliability of cortical activity during natural stimulation. *Trends in cognitive sciences* 14, 40–48 (2010).; Bartels, A. & Zeki, S. Functional brain mapping during free viewing of natural scenes. *Human brain mapping* 21, 75–85 (2004). |

\* The listed references suggest further reading regarding research questions that can be asked using each paradigm, which is not necessarily used in the cited studies. See Langer et al. (2017) for the precise details of all paradigms run as part of this initiative.
**Adapted from:** Langer N et al. (2017) A resource for assessing information processing in the developing brain using EEG and eye tracking *Sci Data*, 4: 170040

*<u>High</u> <u>Density</u> <u>EEG.</u>* High-density EEG data are recorded in a sound-shielded room at a sampling rate of 500 Hz with a bandpass of 0.1 to 100 Hz, using a 128-channel EEG geodesic hydrocel system by Electrical Geodesics Inc. (EGI). The recording reference is at Cz (vertex of the head). For each participant, head circumference is measured and an appropriately sized EEG net is selected. The impedance of each electrode is checked prior to recording to ensure good contact, and is kept below 40 kOhm. Time to prepare the EEG net is no more than 30 min. Impedance is tested every 30 min of recording and saline added if needed.

*<u>Eye</u> <u>tracking.</u>* During EEG recordings, eye position and pupil dilation are also recorded with an infrared video-based eye tracker (iView-X Red-m, SensoMotoric Instruments [SMI] GmbH) at a sampling rate of 120 Hz. This system has a spatial resolution of 0.1° and a gaze position accuracy of 0.5°. The eye tracker is calibrated with a 5-point grid before each paradigm. Specifically, participants are asked to direct their gaze in turn to a dot presented at each of 5 locations (center and four corners of the display) in a random order. In a validation step, the calibration is repeated until the error between two measurements at any point is less than 2°, or the average error for all points is less than 1°.

#### 2.4.6 Magnetic Resonance Imaging (MRI)

*Test Phase (mobile 1.5T Siemens Avanto; n=343).* Imaging data were collected using a 1.5T Siemens Avanto system equipped with 45 mT/m gradients in a mobile trailer (Medical Coaches, Oneonta, NY). The scanner was selected to pilot the feasibility of using a mobile MRI platform to achieve a single scanner solution for the challenges of scanning at geographically distinct locations in the NY area. To maximize long term stability, the trailer was parked on 10-inch thick concrete pads. The system was upgraded with 32 RF receive channels, the Siemens 32-channel head coil, and the University of Minnesota Center for Magnetic Resonance Research (CMRR) simultaneous multi-slice echo planar imaging sequence^31^. Scanning included resting state fMRI, diffusion kurtosis imaging (DKI) structural MRI (T1, T2-space), magnetization transfer imaging, quantitative T1 and T2 mapping (DESPOT T1/T2^32^) and imaging of visceral fat (T1W). See Table 7 for the full scan protocol and Table 8 for parameters.

**Table 7.** MRI Protocol Layout.

| Staten Island |  | Rutgers University |  |
| --- | --- | --- | --- |
| Scan Type | Time (min) | Scan Type | Time (min) |
| Abdomen localizer | 0.52 | Localizer | 0.2 |
| T2Flair | 2.73 | T2Flair | 2.4 |
| Breathhold | 0.18 | fMRI Distortion map | 0.1 |
| Brain localizer | 0.43 | fMRI Distortion map | 0.1 |
| motion training | 1.58 | Rest | 5.1 |
| field map | 1.08 | Peer 1 | 1.9 |
| Resting state | 10.3 | Rest | 5.1 |
| T1W | 6.53 | Peer 2 | 1.9 |
| DWI B=0 PA-AX | 0.27 | Movie: Despicable Me | 10 |
| DKI 64 Directions AP | 9.98 | T1W | 7 |
| DWI B=0 PA-AX | 0.27 | T2Space | 7 |
| DWI B=0 AP-AX | 0.27 | Peer 3 | 1.9 |
| Despot 1 | 5 | Movie: The Present | 4 |
| IR SPRG | 0.88 | MT On | 4 |
| Despot 2 | 5 | MT Off | 4 |
| MT Off | 6.68 | DKI | 10 |
| MT On | 6.68 |  | 64.7 |
|  | 58.4 |  |  |

*Deployment Phase I (3.0T Siemens Tim Trio; ongoing).* Imaging data are collected using a Siemens 3T Tim Trio MRI scanner located at the Rutgers University Brain Imaging Center (RUBIC). The scanner was selected based on physical proximity to the HBN Diagnostic Research Center in Staten Island, New York (12.7 miles; average ride duration: 24 minutes). The system is equipped with a Siemens 32-channel head coil and the CMRR simultaneous multi-slice echo planar imaging sequence. When possible, the structural and functional MRI scan parameters were selected to facilitate harmonization with the recently launched NIH ABCD Study (this was not possible for the diffusion imaging due to limitations of the Trio platform). See Table 7 for scan protocol layout and Table 8 for parameters. Of note, two naturalistic viewing fMRI scans were added to the protocol (“Despicable Me” [10 minute clip; added October 28, 2016], “The Present” [∼4 minutes; added November 23, 2016]).

**Table 8.** MRI Protocol Parameters.

**Staten Island**
|  | Slices | %FOV phase | Resolution(mm) | TR (ms) | TE (ms) | Ti (ms) | Flip Angle (°) | Multi Band Accel | Phase Partial Fourier | Notes |
| --- | --- | --- | --- | --- | --- | --- | --- | --- | --- | --- |
| **T1 MPRAGE** | 176 | 100% | 1.0 × 1.0 × 1.0 | 2730 | 1.64 | 1000 | 7 | N/A | Off |  |
| **T2 FLAIR** | 24 | 87.50% | 0.9 × 0.9 × 5.0 | 9000 | 89.00 | 2500 | 150 | N/A | Off |  |
| **Diffusion** | 72 | 100% | 2.0 × 2.0 × 2.0 | 3110 | 76.20 | N/A | 90 | 3 | 6/8 | 64 directions, b = 0,1000,2000 |
| **fMRI** | 54 | 100% | 2.5 × 2.5 × 2.5 | 1450 | 40.00 | N/A | 55 | 3 | Off |  |
| **MTI** | 176 | 100% | 1.0 × 1.0 × 1.0 | 30 | 11.00 | N/A | 15 | N/A | 6/8 | Acquired with and without MT |

**Rutgers University**
|  | Slices | %FOV phase | Resolution(mm) | TR (ms) | TE (ms) | Ti (ms) | Flip Angle (°) | Multi Band Accel | Phase Partial Fourier | Notes |
| --- | --- | --- | --- | --- | --- | --- | --- | --- | --- | --- |
| **T1 MPRAGE** | 224 | 100% | 0.8 × 0.8 × 0.8 | 2500 | 3.15 | 1060 | 8 | N/A | Off |  |
| **T2 FLAIR** | 22 | 87.50% | 0.9 × 0.9 × 5.0 | 9000 | 90.00 | 2500 | 150 | N/A | Off |  |
| **T2 SPACE** | 224 | 100.00% | 0.8 × 0.8 × 0.8 | 3200 | 564.00 | N/A | varies | N/A | Off |  |
| **Diffusion** | 72 | 100% | 1.8 × 1.8 × 1.8 | 3320 | 100.20 | N/A | 90 | 3 | Off | 64 directions, b = 0,1000,2000 |
| **fMRI** | 60 | 100% | 2.4 × 2.4 × 2.4 | 800 | 30.00 | N/A | 31 | 6 | Off |  |
| **MTI** | 176 | 100% | 1.0 × 1.0 × 1.0 | 30 | 11.00 | N/A | 15 | N/A | 6/8 | Acquired with and without MT |

*Deployment Phase II (3T Siemens Prisma; pending).* In late 2017, Phase II scanning will begin using Prisma scanners located at the CitiGroup Cornell Brain Imaging Center and the CUNY Advanced Science Research Center. The imaging sequence protocols will be harmonized to the NIH ABCD Study.

*Monitoring eye gaze direction during naturalistic viewing.* A key question that may arise in the analysis of naturalistic fMRI scans (or even resting state scans), is where an individual is looking during each repetition in the time series. To address this question, the HBN has included calibration scans that can be used to extract information about eye gaze direction using predictive eye estimation regression (PEER) - a method that uses multivariate regression on calibration data to learn classifiers for decoding eye gaze location from separately acquired fMRI data^33,34,35^. Calibration data involves short fMRI tasks during which the participant is asked to fixate on a white dot that moves through 27 unique positions on the screen. The dot dwells at each location for 4 seconds before moving to a different location and the positions are iterated through twice per calibration run. Two calibration runs are collected during each session, interdigitated with other functional scans, to improve the quality of eye tracking. Once motion corrected, classifiers can be learned from the calibration scans using the voxels in and around the eyes as features and either the x or y coordinate of the dot position as labels. Once learned, the classifiers can be applied to fMRI data to determine where the participant’s eyes were focused during each repetition.

#### 2.4.7 Voice and Video Recording and Actigraphy Data Collection

Behavior monitoring technologies have the potential to help infer internal states of participants during assessments^36^. Voice analysis stands out as particularly promising, given its increasing application in psychiatry (e.g., to assess mood and anxiety^37^), in neurology (e.g., to assess motor function in populations such as those affected by Parkinson’s diseas^38^) and in developmental studies (e.g., to assess pubertal stage^39^). The ease with which one can record audio samples in a controlled setting is particularly appealing. Among sensor-based wearable devices, accelerometer-based actigraphy is a promising means of monitoring behavior related to movement and sleep^40^. For participants in Phase III, the collection of audio and video recordings have begun and actigraphy data collection will be implemented in July 2017.

*<u>Voice recording.</u>* During the administration of all assessments and interviews, starting with subject 746, audio recordings are being collected using a portable Sony ICD-UX 533 digital voice recorder. Additional voice recordings are collected following the MRI Scans. While in the MRI scanner, participants view an animated emotionally evocative four-minute film, titled “The Present”. Immediately after coming out of the scanner, participants are prompted to narrate the story in their own words and answer a series of perspective-taking questions that are related to the film content. During this narration and question answering session, high-fidelity audio recordings are collected with a Røde NT1 cardioid condenser microphone. Additionally, high-definition video of their face and upper body is collected simultaneously with a Canon XC15 digital camcorder. The audio recordings enable voice and speech analysis and the video recordings are envisioned to be useful for facial expression analyses.

*<u>Actigraphy.</u>* Plans are underway to provide each participant with a wrist-worn ActiGraph wGT3X-BT to monitor movement throughout the day and night. Participants will be requested to wear the device every day for one month. The device will be recharged and its data downloaded during each visit.

#### 2.4.8 Genetics

Since December 2016, all participants are asked to provide a saliva sample for genetics using the Oragene Discover (OGR-500) DNA collection kit. This collection strategy was put in place as an alternative to blood collection, which was initially to be carried out in the diagnostic research center, was discontinued due to the logistical challenges it created in the office in the office. Starting in July 2017, saliva samples will be complemented by blood collected in the participant’s home by a local phlebotomy service that the HBN has contracted. Resulting materials will be donated to the NIMH Genetics Repository for sharing.

#### 2.4.9 Deciduous “Baby” Teeth Collection

Beginning in August 2017, all age appropriate participants will be asked to provide 1-2 deciduous “baby” teeth. Shed teeth are collected into provided plastic tubes, labeled, and stored at room temperature until analysis. Teeth will be used to assess environmental exposures throughout the prenatal and early childhood developmental windows. Between 6-13 years of age, children naturally shed 20 deciduous teeth. These teeth begin developing prenatally ∼14-16 weeks after fertilization and mineralization follows a regular incremental layer-by-layer pattern corresponding to the circadian growth rhythm^41^. During development, these layers act as a repository where many chemicals accumulate and offer the opportunity to elicit temporal exposure information. Using bio-imaging along with laser ablation-inductively coupled plasma-mass spectroscopy, researchers leverage the physiology of deciduous teeth to study the intensity and dose of chemical exposure uptake during the pre- and postnatal periods of development^42^. Methods to quantify toxicant and nutrient metal exposures including lead and manganese have been extensively validated^43,44^ and methods are currently under development for a suite of chemical contaminants including additional metals, organic compounds (i.e., phthalates), pesticides, and markers of second hand tobacco smoke and licit/illicit drugs^45^. Additionally, biological markers are being developed for indicators of stress^46^, fetal inflammation, and neurodevelopmental plasticity.

#### 2.4.10 Lessons Learned

Over the course of the development and the implementation of Phases I and II, we have overcome challenges and learned a variety of lessons. Some of the key challenges and solutions that may benefit others are highlighted below:

*1. Incentivizing Participation.* Recruitment is a key challenge for large-scale data generation initiatives, especially when data capture is not simply an add-on to ongoing activities (e.g., addition of a blood sample in clinics or a questionnaire in schools). While scientists commonly justify the funding of research based on potential long-term scientific benefits, the general public tends to evaluate the utility of research participation based on more immediate needs - particularly when participation demands a substantial amount of time and energy. Early in the development of HBN, these competing agendas were repeatedly highlighted by potential community partners. As a result, the HBN has attempted to maximize the quality and breadth of feedback and recommendations that are provided to families and caregivers; the information provided is derived from clinically relevant data obtained over the course of participation (e.g., feedback report and sessions provided by licensed clinicians, generation of a referral grid for the NYC area). From this project’s inception, there has been an emphasis on the distinction between the data obtained purely for research purposes (e.g., EEG, MRI) and the data that may directly benefit participants. This distinction has helped to manage expectations and answer participants’ and family members’ questions about the scope and utility of the project.

*2. Balancing Experimental Needs and Participant Burden.* Drawing from prior experiences with the NKI-Rockland Sample initiative, the HBN was initially designed to be completed in two 6-hour days. Over the course of Phase I, we learned that many participants and their families preferred an alternative schedule that is better aligned with school and work schedules. As a result, the HBN adopted the current schedule of four 3- to 3.5-hour sessions. Despite initial concerns that this would lead to an increased incomplete participation rate, the current schedule has facilitated participation and the dropout rate has remained low at around 6%.

*3. Broadening the Scope of Phenotyping for the Study of Mental Health.* There is a need to consider broader domains of impairment known to be highly associated with psychiatric illness. We received feedback specifically about the desirability of measuring intelligence, learning, language, speech and lifestyle variables (e.g., fitness, eating behaviors, nutrition); in response, we replaced the abbreviated batteries commonly used for intelligence and achievement testing in research studies (e.g., WASI^47^, limited portions of the WIAT) with more comprehensive evaluations (i.e., WISC^16^, full WIAT^48^), which require an additional 90-120 minutes per assessment. In addition to the scientific benefits of expanding the granularity of our evaluations, these evaluations have sometimes been useful for obtaining individualized educational plans (IEPs) for students in the NYC area. Similarly, the addition of screening evaluations for speech and language (followed by more comprehensive evaluations when indicated) resulted in the identification of possible speech or language impairments in 30.6% of the children seen to date.

*4. Logistical Challenges Related to Mobile Data Acquisition.* In part, Phase I was designed to assess the added value of mobile assessment vehicles for data acquisition. In particular, we tested the utility of an MRI scanner housed in a trailer that could be moved periodically (e.g., monthly), as well as a converted mobile recreational vehicle (RV) that was equipped to carry out all non-MRI portions of the assessment. Despite the initial substantial appeal of using these vehicles, logistical issues turned out to be too great. For the mobile MRI scanner, the cost of moving the vehicle more than once a month turned out to be substantial. Even more difficult was finding times to accommodate all eligible participants in the fixed available assessment blocks when the scanner was on-site. Potential data loss when patients were required to wait for the scanner to arrive was also a concern. With regard to the mobile RV for non-MRI assessments, the vehicle worked satisfactorily for staff and participants; however, its throughput was substantially less than what could be obtained in a fixed office space, where multiple participants can be seen simultaneously. Despite the limitations of using the mobile RV for conducting complete evaluations, the vehicle has been a highly effective recruitment tool. Specifically, at community health fairs and events, the vehicle has been used to increase awareness of the project, and to provide short mental health screenings.

*5. Expanding Landscape for Biomarker Identification.* As biomedical and mobile technologies and analysis methods continue to advance, the potential grows for tailored, precise, and accurate digital phenotyping and biomarker identification. Ancillary data consisting of speech samples (audio recordings) and remote movement (actigraphy) have been recently added to the HBN assessment protocol. We are evaluating other wearable devices with sensors that track physiological state, such as electrodermal activity to monitor stress and photoplethysmography to monitor heart rate. Collection of hair samples (for determination of current metal levels) and of baby teeth (for determining fetal exposure to various metals^49^) are being added to the protocol. Microbiomics is a potentially valuable avenue of exploration that is gaining increased attention, but fecal and other microbiome data collection are being deferred until the practical considerations that such data collection entail can be worked out.

*6. Balancing Efficiency, Innovation and Tolerability of MRI Scan Protocols.* Maximizing tolerability of the scanner environment and minimizing head motion are two inherent challenges for MRI studies, particularly those focusing on developing and clinical populations. Consistent with its predecessor initiative, the NKI-Rockland Sample^12^, the HBN initially included a 10-minute resting state scan. However, head motion was found to be problematic, particularly in the second half of the scan. To address this concern, the resting state scan was eventually broken into two 5-minute scans at the Rutgers data collection site, and removed altogether for 5 year olds, where data quality concerns were most notable. Additionally, for the deployment phase, experimental structural images (e.g., quantitative T1/T2 mapping) were removed in favor of increasing functional MRI scan time. Rather than adding more resting state fMRI scans, we opted to add naturalistic viewing (i.e., movie watching) fMRI sessions to reduce motion and to permit a broader range of analyses^50,51^.

*7. Inclusion of Consent for Commercial Use.* The research community increasingly aims to generate data and methods that will form the foundation of clinically useful tools. As the field attempts to market and distribute innovations, there will be a growing need for the involvement of commercial entities. In preparation for this next phase, we followed NIH recommendations and integrated a consent document for commercial use into the informed consent (starting with participant #701). The receipt of such permission is essential to avoid any ethical or legal concerns that may arise from the commercial use of data for participants who did not provide explicit permission.

8. *Extension of Questionnaire Age Ranges.* Initially, for each questionnaire, determination of whether to administer it to all participants or to a select age group was based on ages indicated by publisher websites, or from validation studies (e.g., ages 8-18 for the SCARED^52^). While this is generally sensible for self-report versions of questionnaires, particularly when reading level is an issue, we have called into question the value of the decision for questionnaires completed by parents. Although some parent-report questionnaires have in the past been used only for ages 8 and up, or up to age 17, this does not mean they cannot be informative for the purposes of the HBN and lack of previously established norms (e.g., t-scores) may be overcome given the magnitude of the data (e.g., the SCARED). Thus, we have reviewed each questionnaire carefully and expanded the age ranges so that parent-report questionnaires are now collected for participants of all ages (5-21) except where developmentally inappropriate (e.g., substance use questionnaires, puberty questionnaires), or where age-specific versions of the same form exist (e.g., ASEBA forms^26^). Increasing the age range for questionnaire administration minimizes data loss in the sample, particularly in the youngest and oldest participants. Additionally, collecting data from broader age ranges may help support extension of normative ranges.

## 3. DATA RECORDS

### 3.1 Data Privacy

During the consent process, all participants provide informed consent for their data to be shared via IRB-approved protocols. Data sharing occurs through the 1000 Functional Connectomes Project and its International Neuroimaging Data-sharing Initiative (FCP/INDI)^53^. Prior to entry of data into the HBN Biobank, all personal identifiers specified by the Health Insurance Portability and Accountability Act (HIPAA) are removed, with the exception of zip code (which is only shared upon request following completion of the HBN Data Usage Agreement described below in section 3.2.1).

Given the sensitive nature of the information provided during HBN participation, a Certificate of Confidentiality was obtained from the Department of Health and Human Services (HHS). The certificate helps to protect the privacy of human subjects by allowing the research team to refuse to disclose names or other identifying characteristics of study participants in response to legal demands (https://humansubjects.nih.gov/coc/index).

### 3.2 Distribution for Use

#### 3.2.1 Phenotypic Data

Phenotypic data may be accessed through the COllaborative Informatics and Neuroimaging Suite (COINS) Data Exchange (http://coins.mrn.org/dx) or an HBN-dedicated instance of the Longitudinal Online Research and Imaging System (LORIS) located at http://data.healthybrainnetwork.org/.

With the exception of age, sex and handedness, which are publicly available with the imaging, EEG and eye-tracking datasets, the HBN phenotypic data are protected by a Data Usage Agreement (DUA). Investigators must complete and have approved by an authorized institutional official before receiving access (the DUA can be found at: http://fcon_1000.projects.nitrc.org/indi/cmi_healthy_brain_network/sharing.html). Modeled after the practices of the NKI-Rockland Sample, the intent of the HBN DUA is to ensure that data users agree to protect participant confidentiality when handling the high dimensional HBN phenotypic data (which includes single item responses), and that they will agree to take the necessary measures to prevent breaches of privacy. With the exception of zip code (which is only available upon request), no protected health identifiers are present in data distributed through the DUA, as a means of ensuring minimal risk of privacy breach. The DUA does not place any constraints on the range of analyses that can be carried out using the shared data, nor does it include requirements for co-authorship by the originators of the HBN Biobank.

#### 3.2.2 EEG, Eye-tracking, and Imaging Data

All EEG, eye tracking and imaging data can be accessed through the 1000 Functional Connectomes Project and its International Neuroimaging Data-sharing Initiative (FCP/INDI) based at http://fcon_1000.projects.nitrc.org/indi/cmi_healthy_brain_network. This website provides an easy-to-use interface with point-and-click download of HBN datasets that have been previously compressed; the site also provides directions for those users who are interested in direct download of the data from an Amazon Simple Storage Service (S3) bucket. Imaging data is stored in the Brain Imaging Data Structure (BIDS) format, which is an increasingly popular approach to describing MRI data in a standard format^54^.

All data are labeled with the participant’s unique identifier. EEG data are available openly, along with basic phenotypic data (age, sex, handedness, completion status of EEG paradigms) and performance measures for the EEG paradigms. These data are located in a comma-separated (.csv) file accessible via the HBN website.

### 3.3 Partial and Missing Data

Some participants may not be able to successfully complete all components of the HBN protocol due to a variety of factors (e.g., participants experiencing claustrophobia may not be able to stay in the scanner for the full session; a participant with sensory issues may have a more limited ability to participate in the EEG protocol). To prevent data loss when possible, we include exposure procedures such as a mock MRI scanner experience during visit 1, and repeat exposures to an EEG cap prior to session 4. Overall, we attempt to collect as much of the data as possible within the allotted data collection intervals and log data losses when they occur.

### 3.4 Data License

HBN imaging, EEG and eye-tracking datasets for the first 701 participants enrolled are currently distributed under the Creative Commons, Attribution Non-Commercial Share Alike 4.0 International Public License (https://creativecommons.org/licenses/by-nc-sa/4.0/), as they were collected prior to the addition of consent for commercial use to the informed consent (specific participant IDs are specified on the HBN data-sharing website). From December 6, 2016 forward, HBN datasets are being distributed using the Creative Commons Attribution 4.0 International Public License (https://creativecommons.org/licenses/by/4.0/), which does allow for commercial use of datasets. For the high-dimensional phenotypic data, all terms specified by the DUA must be complied with.

## 4. TECHNICAL VALIDATION

### 4.1 Quality Assessment

Consistent with policies established through our prior data generation and sharing initiatives (i.e., FCP/INDI^53^; NKI-Rockland Sample^12^), all imaging datasets collected through the HBN are being made available to users, regardless of data quality. This decision is justified by a lack of consensus in the imaging community on what constitutes “good” or “poor” quality data. Also, “lower quality” datasets can facilitate the development of artifact correction techniques and of evaluating the impact of such real-world confounds on reliability and reproducibility. Given the range of clinical presentations in the HBN, the inclusion of datasets of varying quality creates a unique opportunity to test for associations with participant-related variables of interest beyond age and hyperactivity (e.g., anxiety, autistic traits).

#### 4.1.2 Phenotypic Data

Beyond checking data for outliers, a key question for the evaluation of phenotypic data is whether or not the observed distributions and inter-relationships are sensible. Figure 2 depicts the distribution of sample variables of interest related to mental health and learning. As can be seen, the data obtained for variables known to have a normal distribution (e.g., IQ) exhibited a normal distribution in the HBN dataset. Of note, the total score from the Child Behavior Checklist, a measure that typically only has meaningful variation among symptomatic individuals (resulting in a truncated distribution), was found to have a broad distribution in the HBN that was close to normal; this represents the wide variation in symptom severity across the range of phenotypic measures which is inherent to the HBN recruitment strategy.

**Figure 2.**
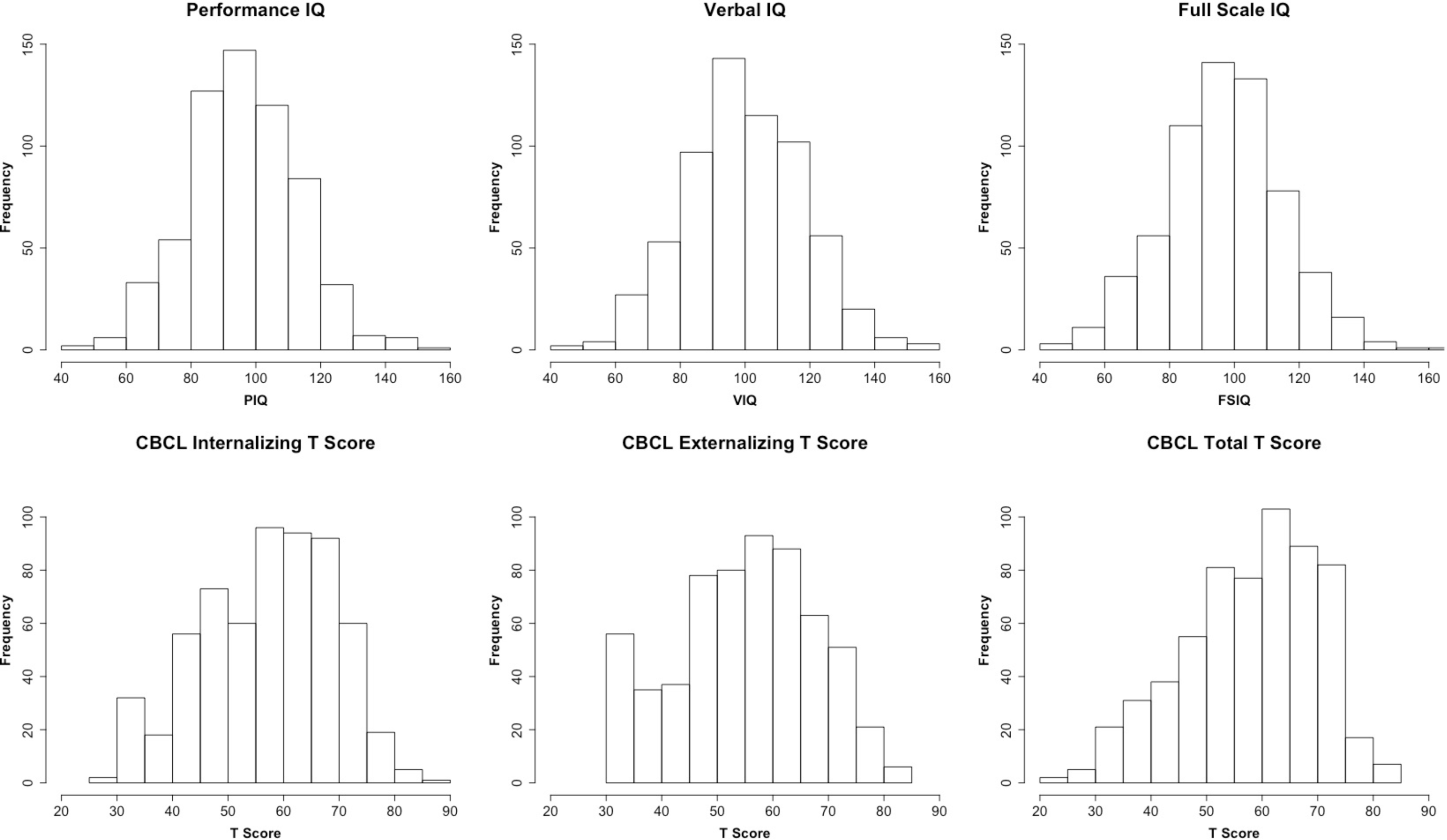
*Distribution of IQ measures and CBCL Scores.* Participant IQ was measured using the WISC, with the exception of: 1) early participants for whom the more abbreviated WASI was performed, 2) individuals with limited verbal skills and/or known IQ less than 70, or 3) children under age 6. For these latter two cases, the KBIT was performed. These figures include overall performance IQ, verbal IQ, and full-scale IQ measures from all three tests.

To further facilitate the evaluation of phenotypic data, we plotted correlations between a broad sampling of measures included in the HBN (see Figure 3). Statistical relationships observed after false discovery rate-based correction for multiple comparisons revealed a wealth of associations that are in general alignment with the broader psychiatric literature. For example: 1) at the most basic level, socioeconomic status (Barratt Simplified Measure of Social Status^55^) was positively associated with indices of intelligence (Full scale IQ [FSIQ], Performance IQ [PIQ], Verbal IQ [VIQ]) and language performance (i.e., CELF screener), and negatively associated with multiple indices of mental illness, 2) general measures of internalizing and externalizing symptoms exhibited high correlations with one another, 3) autistic and ADHD traits were each negatively associated with performance on intelligence tests, 4) prosocial tendencies were higher in those with lower levels of symptoms related to ADHD traits, autistic traits and affective reactivity, 5) higher body mass index was associated with internalizing symptoms and increased peer problems. Of note, parent report for anxiety appeared to reveal more robust relations with other measures (e.g., autistic traits) than did child self-report, consistent with expected rater-bias effects.

**Figure 3.**
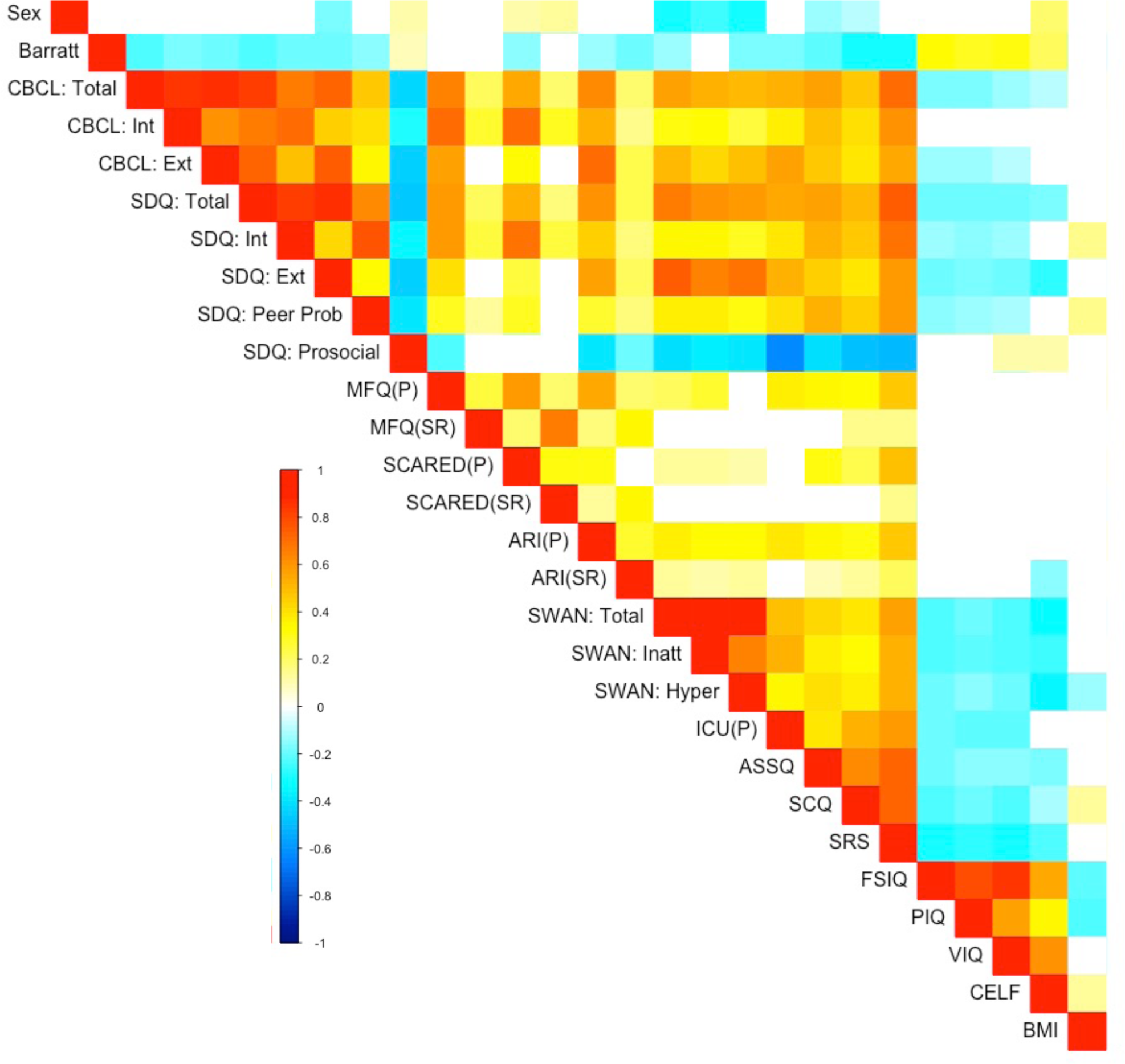
*Correlation Matrix of HBN Phenotypic Measures:* Heatmap depicts significant correlations between a broad sampling of HBN behavioral, cognitive, and physical measures after multiple comparisons correction (false discovery rate; q < 0.05). The associations revealed are in general alignment with the broader psychiatric literature.

#### 4.1.3 Imaging Data

Consistent with recent major FCP/INDI data releases (i.e., the Consortium for Reliability and Reproducibility [CoRR]^56^, Autism Brain Imaging Data Exchange 2 [ABIDE2]^57^), we made use of the Preprocessed Connectome Project Quality Assurance Protocol (QAP)^58^ to assess data quality for core MRI data modalities (i.e., functional MRI, morphometry MRI and diffusion MRI). The QAP includes a broad range of quantitative measures that have been proposed for assessing image data quality (see Table 9 for list of measures and their definitions, adapted from^57^).

**Table 9.** Description of the Preprocessed Connectome Project (PCP) Quality Assurance Protocol (QAP) Measures.

| Spatial Metrics | Description |
| --- | --- |
| Contrast-to-noise ratio (CNR)<sup>85</sup> (sMRI only) | $M_{GM} \text{ intensity} - M_{WM} \text{ intensity} / SD_{air} \text{ intensity}$. *Larger values reflect a better WM GM distinction.* |
| Signal-to-noise ratio (SNR)<sup>85</sup> | $M_{GM} \text{ intensity} / SD_{air} \text{ intensity}$. *Larger values reflect less noise* |
| Artifactual voxel detection (Qi1)<sup>86</sup> (sMRI only) | * voxels with intensity corrupted by artifacts/ *voxels in the background. *Larger values reflect more artifacts which likely due to motion or image instability.* |
| Entropy Focus Criteria (EFC)<sup>87</sup> † | Shannon's entropy of each voxel's intensity used to measure ghosting and blurring due to head motion. *Larger values reflect more blurring likely to motion or technical differences.* |
| Smoothness of Voxels (FWHM)<sup>88</sup> † | Full-width half maximum of the spatial distribution of the image intensity values. *Larger values reflect more spatial smoothing maybe due to motion or technical differences.* |
| Foreground to Background Energy Ratio (FBER) † | M energy of image intensity (i.e., mean of squares) within the head relative to that of outside the head. *Larger values reflect higher signal in relation to noise.* |
| Ghost to Signal Ratio (GSR)<sup>89</sup> † | M signal in the 'ghost' image divided by the M signal within the brain. *Larger values reflect more ghosting likely due to physiological noise, motion, or technical issues.* |
| Temporal Metrics (fMRI * and DTI only) | Description |
| Mean framewise displacement- Jenkinson (mFD)<sup>59</sup> ‡ | Sum absolute displacement changes in the x, y and z directions and rotational changes around them. Rotational changes are given distance values based on changes across the surface of a 50 mm radius sphere. *Larger values reflect more movement.* |
| % and * volumes with FD>0.2 mm ‡ | % and *volume to volume motion >0.2 mm FD. *Larger values reflect more movement.* |
| Standardized DVARS<sup>90</sup> ‡ | Spatial SD of the data temporal derivative normalized by the temporal SD and autocorrelation. *Larger values reflect larger frame-to-frame differences in signal intensity due to head motion or scanner instability.* |
| Outlier Detection<sup>91</sup> † | M fraction of outliers in each volume per 3dToutcount AFNI command. *Higher values reflect more outlying voxels, which may be due to scanner instability or RF artifacts.* |
| Global Correlation (GCORR) ‡ | M correlation of all combinations of voxels in a time series. Illustrates differences between data due to motion/physiological noise. *Larger values reflect a greater degree of spatial correlation between slices, which may be due to head motion or 'signal leakage' in simultaneous multi-slice acquisitions.* |
| Median Distance Index<sup>91</sup> ‡ | M distance (1—spearman's rho) between each time-point's volume and the median volume using AFNI's 3dTqual command. *Higher values reflect greater differences between subsequent frames, which may be due to head motion or technical issues.* |

\* For all R-fMRI data temporal metrics have been computed after discarding the first 5 time points of the time series which were field map corrected if field
† For R-fMRI data these metrics are computed on mean functional data.
‡ For R-fMRI these metrics are computed on time series data. M, Mean; GM, Gray Matter; WM, White Matter; s.d., Standard Deviation.
**Adopted from** Di Martino A, et al. 2017. Enhancing studies of the connectome in autism using the autism brain imaging data exchange II. Sci Data. 4:170010.

Given commonly cited concerns about head motion during MRI scans, particularly during resting state fMRI scans, we examined age-related differences in motion. We quantified head motion using frame-wise displacement (FD), which is calculated using root mean square deviation^59^. Mean FD is commonly used to evaluate the impact of movement on a dataset^60,61^, but it cannot distinguish between occasional large movements and frequent smaller movements, the effects of the former being likely easier to fix using motion scrubbing or volume censoring methods^60^. Consistent with this concern, Figure 4 panel A demonstrates a nonlinear relationship between mean FD and median FD, with the latter providing a better indication of the amount of the data that can be retained after movement correction (e.g., volume censoring).

**Figure 4.**
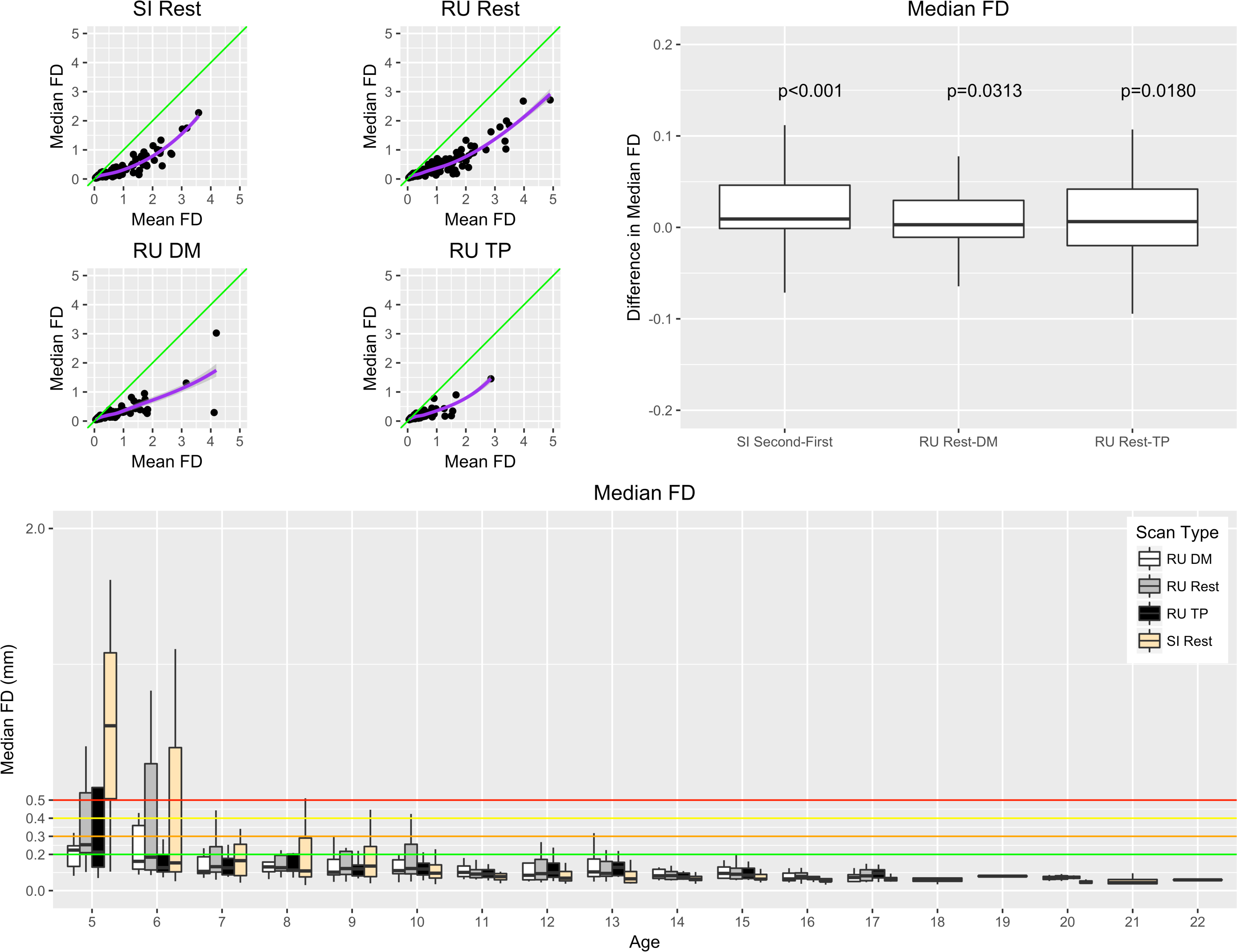
*Median Framewise Displacement Measures.* The upper left panel plots Median Framewise Displacement (Median FD) vs. Mean Framewise Displacement (FD) for Staten Island (SI) and Rutgers (RU) fMRI. The upper right panel shows the difference in median FD between different scan conditions. The bottom panel shows median FD for different scan types for different ages. Significance values depicted reflect results of paired t-test.

Consistent with prior work^62^, both sites (the 1.5 Tesla mobile scanner in Staten Island and the 3.0 Tesla fixed scanner at Rutgers University) exhibited negative associations between age and head motion for all functional scan types, with children between ages 5 and 8 exhibiting the greatest levels of movement. Median FD tended to be higher during the second half (5 minutes) of the resting state scan than during the first half; this observation resulted in our decision to split the scan into two 5-minute scans starting with participant 538 in February 2017. As predicted by recent work highlighting the advantages of naturalistic viewing to minimize head motion, we found that head motion was significantly reduced during each of the movie-watching scan sessions (“Despicable Me” [n = 307], “The Present” [n = 251]) relative to rest.

Beyond the examination of temporal characteristics of the HBN data, we also applied the structural measures included in the PCP QA to each of the core data types (functional, diffusion, morphometry). See Figure 5 for a subset of these measures; the full set of measures are included on the HBN website in a comma-separated tabular format for download.

**Figure 5.**
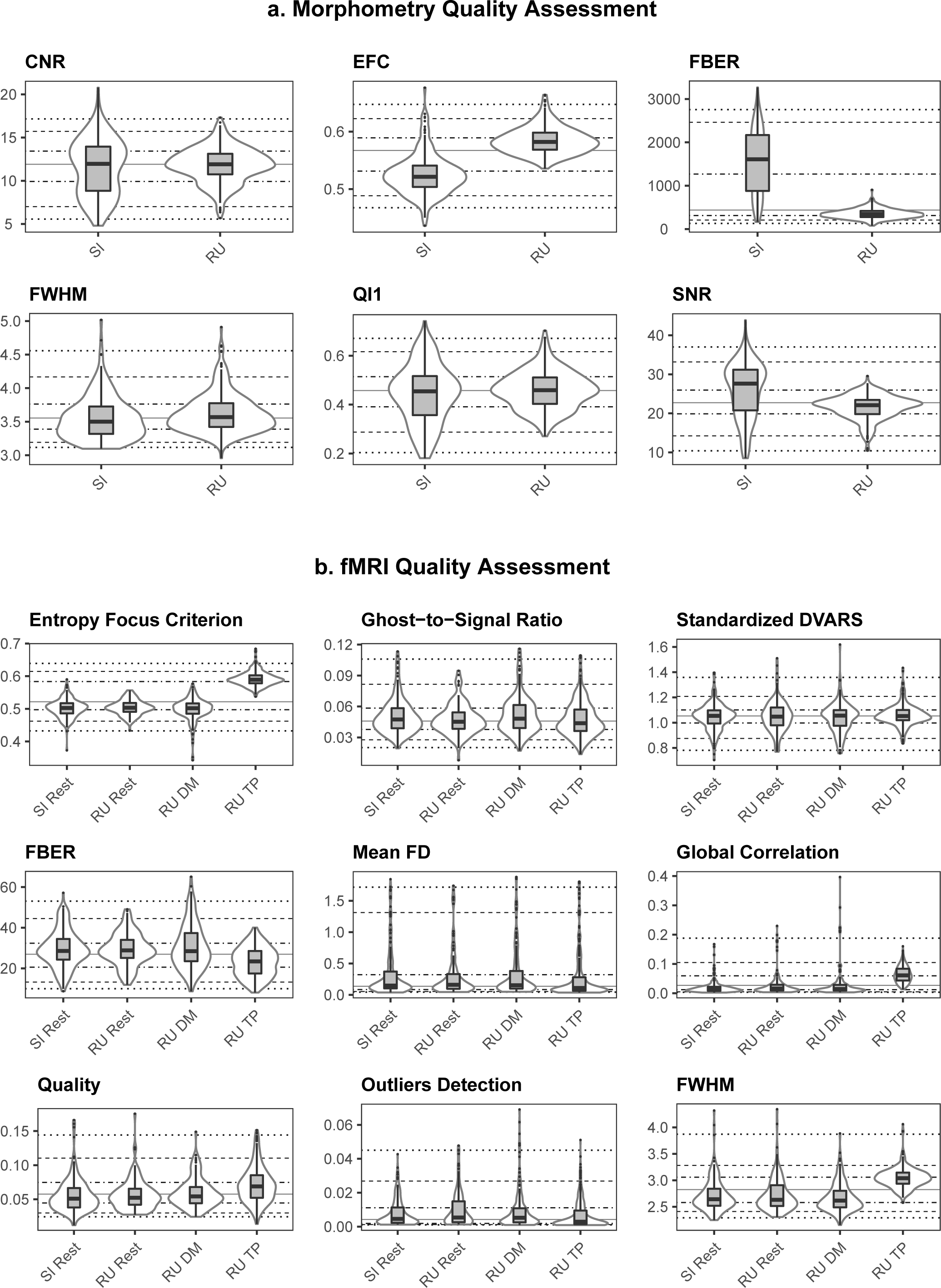
*Preprocessed Connectome Project Quality Assurance Measures for functional and morphometric MRI.* Shown here are PCP QA results for morphometry (upper panel) and functional (lower panel) MRI data quality for each data acquisition phase - Staten Island (SI; 1.5 Tesla Siemens Avanto) and Rutgers (RU; 3.0T Siemens Tim Trio).

#### 4.1.4 Associations Between Imaging QA and Clinical Variables

With the range of clinical presentations and ages present in the HBN, there is a unique opportunity to test for associations between phenotypes and dimensions of data quality. Figure 6 depicts significant relationships detected between phenotypic variables and QAP parameters for the different scans, using Pearson correlation (only significant relationships, surviving false discovery rate correction for multiple comparisons, are depicted). Not surprisingly, for fMRI, age was negatively associated with nearly all motion indices, regardless of scan type. Interestingly, while motion parameters were correlated with an ADHD measure of hyperactivity during the rest condition, they did not correlate significantly during the movie conditions; these findings are in accord with the suggestions of prior work examining the impact of movies on head motion^63^. The quality assurance associations with behavioral variables of interest highlighted here are not intended to be dissuasive, but rather to emphasize the importance of considering and accounting for the potential contributions of data quality to higher order analyses.

**Figure 6.**
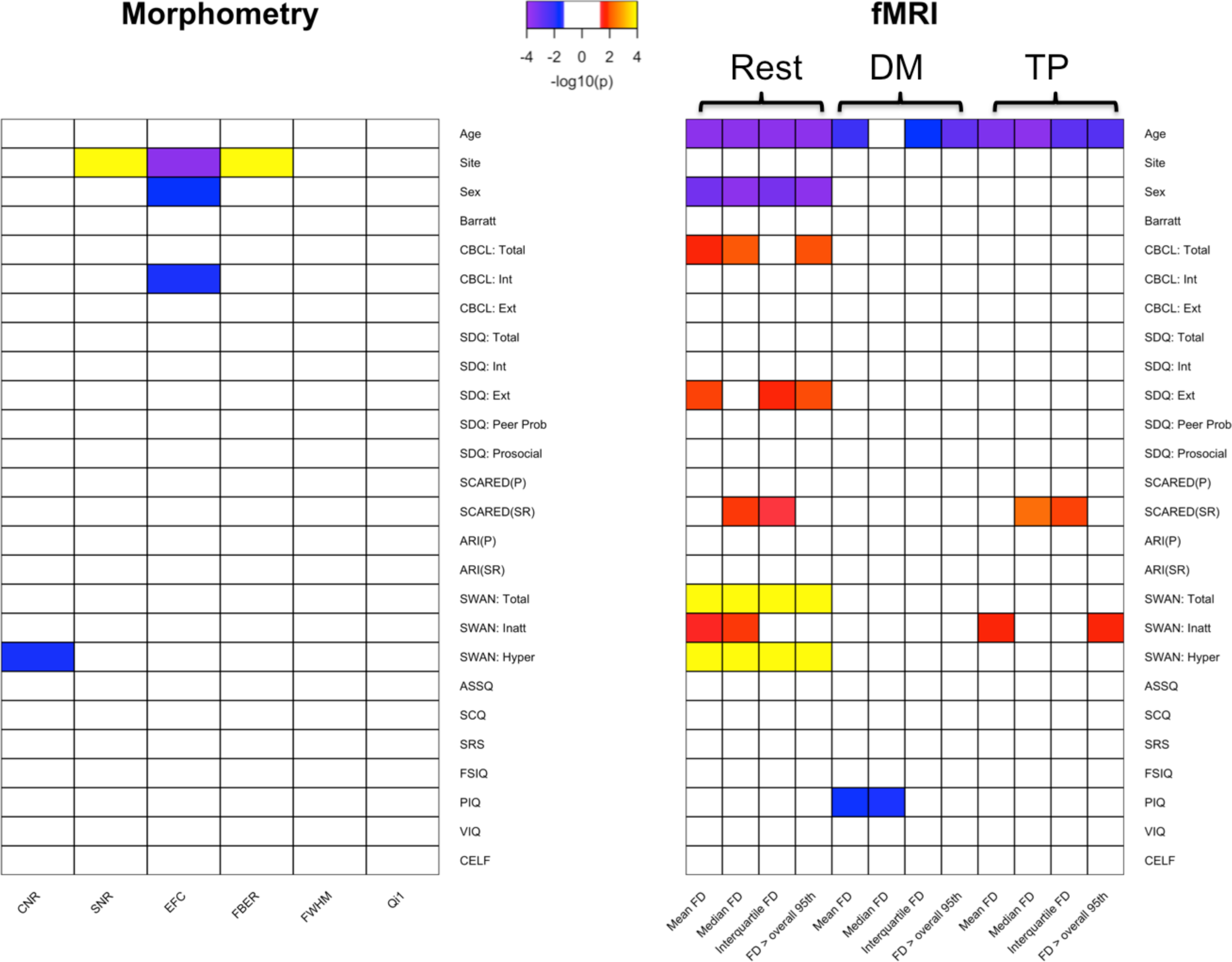
*Correlation Between Phenotypic Measures and QAP measures.* Here we depict significant Pearson correlations (after false discovery rate correction for multiple comparisons) between phenotypic measures and key QA indices for morphometry MRI (left panel), as well as each of the functional MRI scan types (resting state fMRI, naturalistic viewing fMRI: ‘Despicable Me’, naturalistic viewing fMRI: ‘The Present’) (right panel). To facilitate visualization, significance values are depicted as -log10(p).

#### 4.1.5 EEG Data

For each of the EEG acquisitions, Figure 7 depicts the number of channels rejected based on the data distribution and variance of channels (threshold: > 3 standard deviations), as implemented in EEGLAB’s *pop_rejchan.m* function^64^.

**Figure 7.**
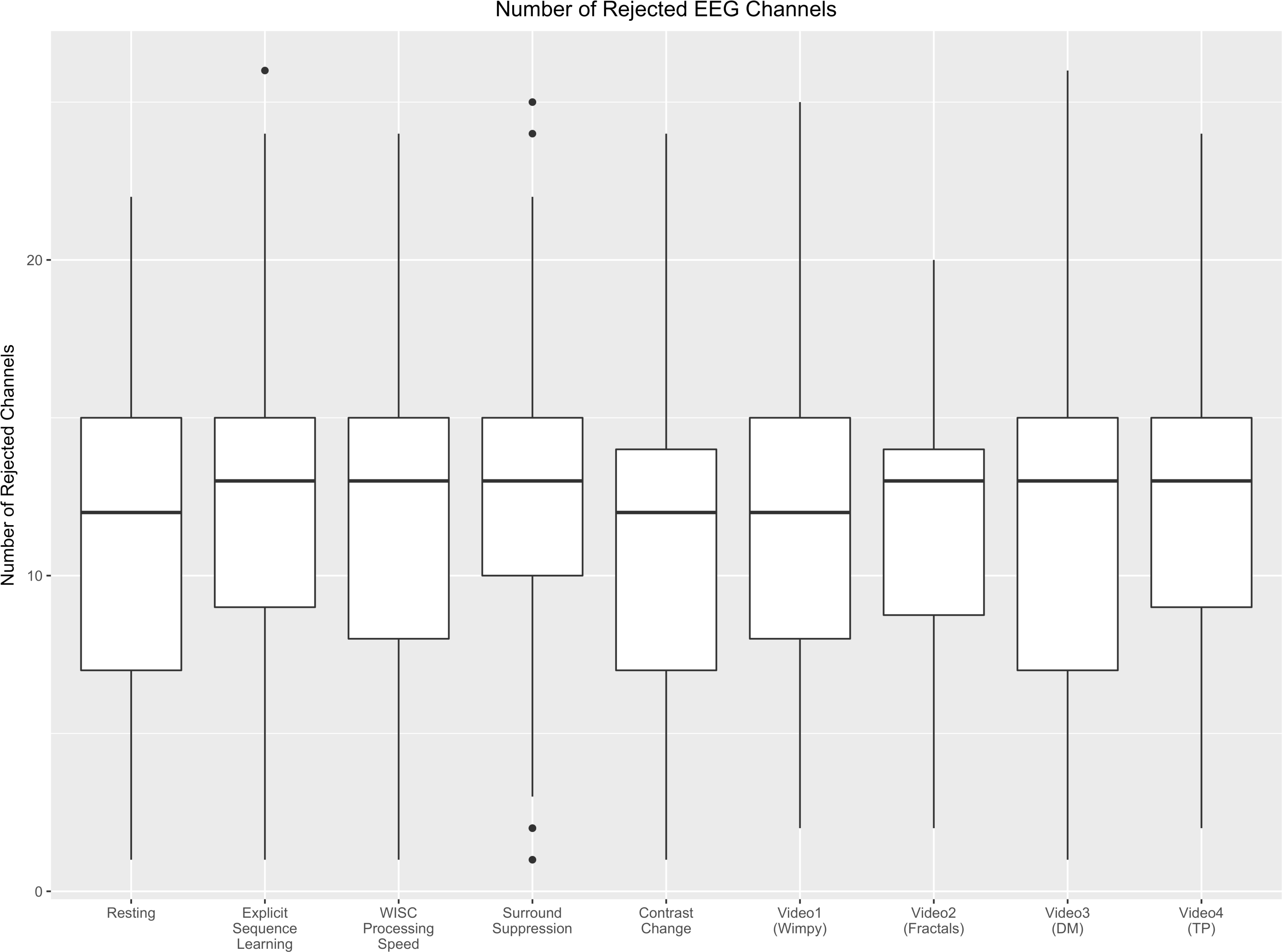
EEG quality assessment: Shown here are the number of rejected EEG channels for each of the paradigms.

#### 4.1.6 Sampling Biases and Representativeness

Although relatively early in recruitment, there is sufficient data to obtain insights into potential biases arising in the HBN sample, as well as its representativeness of the general population. One of the most notable biases is the over-representation of males relative to females in the first release (2:1) (Figure 8). A few factors may account for this. First is the prevalence of ADHD in the sample, a disorder that is commonly estimated to have 3:1 male:female ratio in children (Figure 9). The prominence of ADHD in the sample is not surprising as it is among the most prevalent childhood disorders, and given that it is an externalizing disorder, it is much less likely to go unnoticed than internalizing disorders (e.g., current estimates suggest that as many as 80% of individuals with anxiety disorders go undiagnosed and untreated)^65^. Another factor contributing to prominence of ADHD may be the current age distribution; median age in the initial release is 10.7 years old, with an interquartile range of 7.8-13.3 (Figure 8). Heavier weighting towards childhood and early adolescence may explain lower rates of internalizing relative to externalizing disorders. Future recruitment will include targeted efforts to increase the representation of internalizing disorders and older adolescents. Similarly, as sample size continues to grow and the additional diagnostic research center intended for Harlem is added, we will monitor community variables (e.g., household income, parental education, parental marital status, and race/ethnicity).

**Figure 8.**
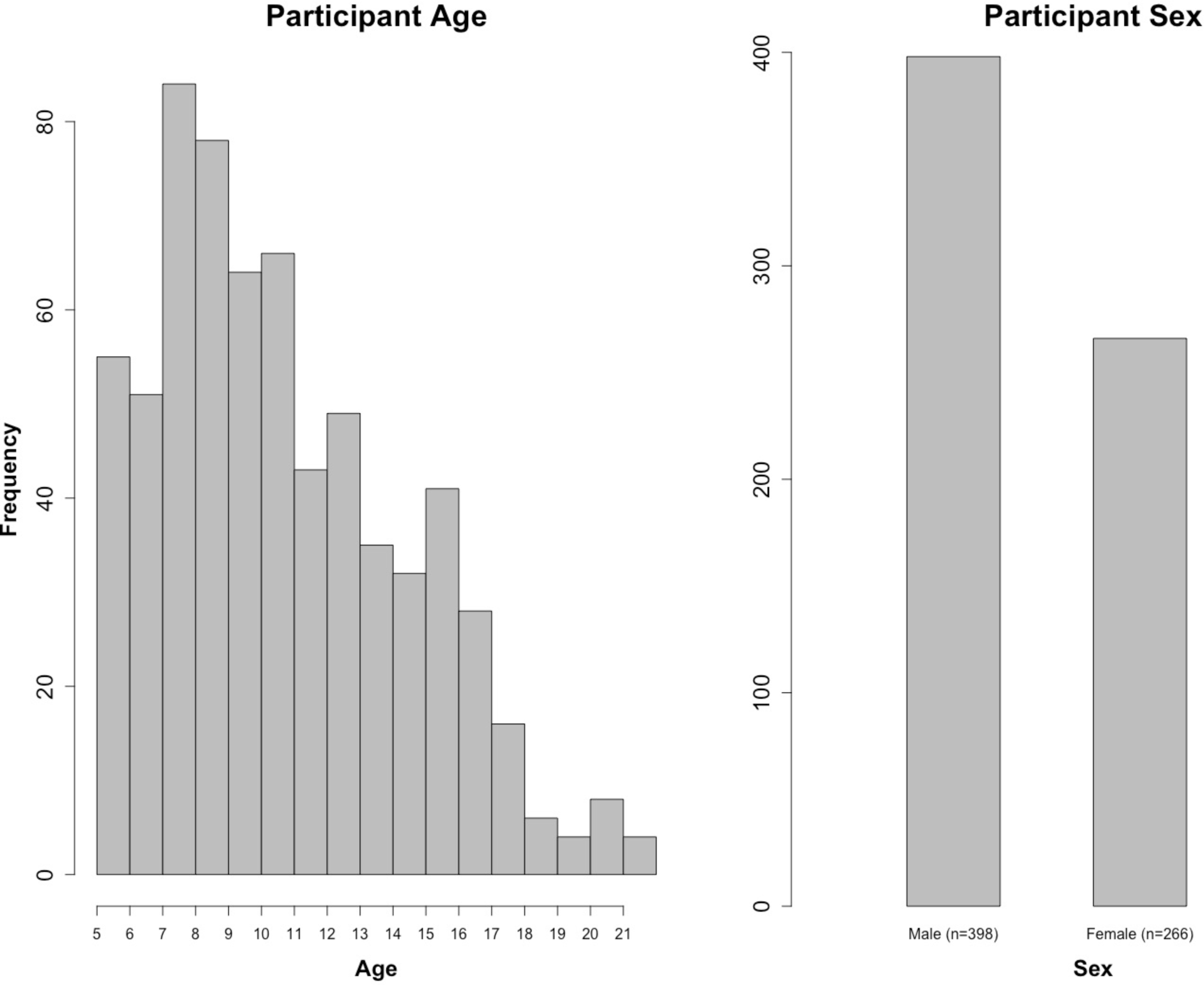
*Age and Sex Distribution of HBN Participants.*

**Figure 9.**
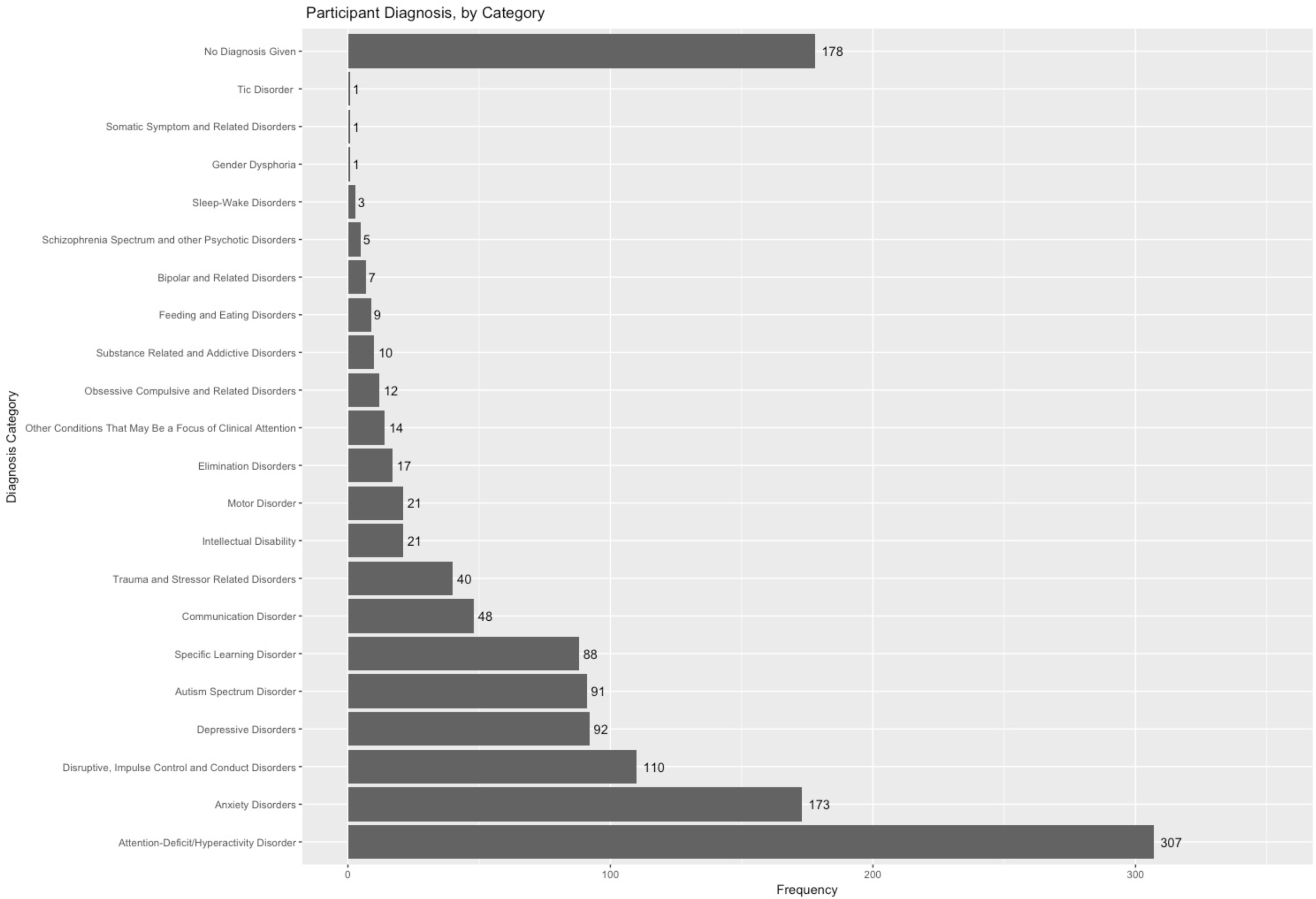
*Diagnostic Breakdown of HBN Participants.* This figure shows the frequency of diagnoses given to HBN participants. Data for this figure comes from the final consensus diagnosis given by the lead clinician at the end of participation. Diagnoses are grouped by category.

## 5. USAGE NOTES

### 5.1 Handling Head Motion in MRI Data

Head motion presents an unavoidable challenge for developmental and clinical imaging, regardless of MRI modality (fMRI, dMRI, sMRI). Arguably, the most basic strategy for handling motion, short of applying an uncomfortable motion-restricting apparatus, is limiting analyses to high-quality data. The Brain Genomic Superstruct data release is an excellent example of the utility of large-scale datasets in supporting such a strategy, as 1,570 datasets were selected for analyses from a pool of 3,000 individuals following rigorous quality control^66^. A limitation of this strategy for psychiatric data is that many phenotypes of interest are inherently more prone to head motion (e.g., children under 9, those with Attention-Deficit/Hyperactivity Disorder), especially those with higher symptoms levels. Compounding the downsides of discarding data are the increased costs associated with the recruitment and phenotyping of clinical populations.

For functional MRI, an alternative strategy is to statistically correct the data for movement-induced intensity fluctuations, or remove offending time frames altogether^60^. This can be accomplished by a number of means, ranging from regressing a model of movement from the data (e.g., spike regression^67^), removing the contributions of motion-related spatial patterns^68^), attenuating motion spikes using a squashing function, removing offending frames, zeroing out offending frames, or deleting offending frames followed by interpolation. More generalized correction approaches, such as global signal regression and forms of white matter and cerebrospinal fluid regression (e.g., tCompCor, aCompCor^69,70^) can also help to account for motion artifacts. While there is no consensus approach to date, there is a growing literature focused on providing benchmark evaluations of these approaches, as well as their relative merits and weaknesses (e.g., see^61, 71^), that can be used to help select among these corrections.

More broadly, group-level statistical corrections can be used to account for the contributions of motion-related artifacts to associations revealed through data analysis^67^. In the case of functional MRI, this can be accomplished by including motion parameters as a statistical covariate at the group level. Given the trait nature of head motion^56^, some have advocated for using fMRI-derived motion parameters in structural analysis as well. Alternatively, accounting for full-brain differences in measures of interest at the group-level has been shown to be a potentially valuable approach to minimizing the deleterious effects of motion, particularly for fMRI^71^.

It is our hope that the breadth of the Healthy Brain Network dataset will provide a practical perspective on the challenges of motion for various domains of illness and help to stimulate continued development and testing of novel correction strategies.

### 5.2 Special Opportunities

The HBN Biobank is intended to be a resource for accelerating the pace of scientific advancement for neurodevelopmental and learning disorders, and accomplishing this goal will require the combined expertise of a wide range of disciplines. From high-performance computing strategies for addressing the scale of the data, to new analytical strategies for performing regressions on graphs, and better instruments for assessing dimensions of cognitive development, there are a variety of ways researchers can use these data. Below are a few research questions that we believe will be particularly suitable for these data.

#### 5.2.1 Advancing Biophysical Modeling (EEG, fMRI, dMRI)

Mathematical models are an increasingly popular tool for establishing links between brain function and structure. Although still early in their development, recent applications have demonstrated the ability of biophysical models to make predictions about patterns of brain function assessed using fMRI and EEG, as well as behavior^72,73^. The inclusion of fMRI, EEG and diffusion imaging in the HBN will help from the data (AROMA investigators to build bridges between these three modalities, as well as the underlying morphology, for which increasingly sophisticated characterizations are being afforded by automated pipelines, such as MindBoggle^74^. Such models can also be useful for developing and testing hypotheses about possible mechanisms underlying variations in behavior, as well as the occurrence of disease states. Additionally, researchers will be able to test the ability to compare the result of EEG-based functional connectivity analyses carried out in source space (i.e., anatomical space following source localization), with those obtained using functional MRI; such comparisons are important for those interested in the development of clinical tools, as EEG is easier to administer and has lower costs.

#### 5.2.2 Naturalistic Viewing EEG and fMRI

A growing literature over the past decade supports naturalistic viewing EEG and functional MRI^75–77^. Akin to the arguments for resting state fMRI methods nearly a decade ago, advocates highlight findings of reliability for various phenomena observed with naturalistic viewing, as well as the potential to assess inter-individual differences^78^. Recent works have suggested that naturalistic viewing may yield equivalent or even superior levels of reliability for the assessment of functional connectivity relative to rest^79,80^, with the potential to yield novel functional connectivity measures (e.g., inter-subject functional connectivity)^50^ (*Note:* The Healthy Brain Network Serial Scanning Initiative is an openly available resource carried out in preparation for the HBN, which was inspired in part by the MyConnectome Project [http://myconnectome.org/wp/^81^] and can be used to carry out comparisons of the reliability and comparability of differing scan states [e.g., rest, naturalistic viewing, task fMRI]). Finally, the naturalistic viewing experimental paradigm can be used for the study of temporal dynamics in the brain^50^. To facilitate the translation of findings between EEG and fMRI, the animated film titled “The Present”, directed by Jacob Frey, is now included in both the HBN EEG and fMRI protocols. To date, 248 participants have watched “The Present” during EEG, 251 participants have watched during fMRI, and 129 participants have watched during both EEG and fMRI.

#### 5.2.3 Questionnaire refinement and applications of item response theory

A key reality for biologically focused studies is that the potential for discovery is limited by the quality and breadth of phenotyping. The breadth of questionnaires and measures in the HBN provides opportunities for deriving optimal measure sets that minimize the number of items required to characterize an individual while maximizing their predictive value. Beyond traditional factor analyses, item response theory^82–84^ is promising to accelerate the process of finding those questions or measures that are most essential for characterizing differences between individuals.

#### 5.2.4 Voice analysis for biomarker identification

Extraction and analysis of high-dimensional feature sets to characterize vocal production, speech patterns, and speech content is a promising direction for biomarker identification. Features characterizing vocal production are independent of speech content itself, and can provide objective measures of motor difficulties as well as independent means of assessing psychiatrically relevant states, such as mood and anxiety. Features related to patterns and content of speech provide additional opportunities to characterize more complex emotional and cognitive states, as well as issues related to processing information and expressing thoughts. Coupled with other behavioral assessments in the HBN protocols, voice and speech data will encourage users of the HBN data to consider richer and more nuanced approaches to analyzing phenotypic data.

## AVAILABILITY OF SUPPORTING DATA

### LIST OF ABBREVIATIONS

ABCD: Adolescent Brain Cognitive Development
ABIDE: Autism Brain Imaging Data Exchange
ADHD: Attention-Deficit/Hyperactivity Disorder
ADOS: Autism Diagnostic Observation Schedule
AROMA: Automatic Removal of Motion Artifacts
BIA: Bioelectrical Impedance Analysis
CELF-5: Clinical Evaluation of Language Fundamentals, 5th Edition
CMI: Child Mind Institute
COINS: COllaborative Informatics and Neuroimaging Suite
CoRR: Consortium for Reliability and Reproducibility
CTOPP: Comprehensive Test of Phonological Processing
CUNY: City University of New York
dMRI: diffusion magnetic resonance imaging
DSM-5: Diagnostic and Statistical Manual of Mental Disorders, 5th Edition
DUA: data usage agreement
EEG: electroencephalogram
EGI: Electrical Geodesics, Inc.
EVT: Expressive Vocabulary Test
FCP: 1000 Functional Connectomes Project
fMRI: functional magnetic resonance imaging
FSIQ: full scale intelligence quotient
GFTA: Goldman Fristoe Test of Articulation
HBN: Healthy Brain Network
HIPAA: Health Insurance Portability and Accountability Act
IEP: Individualized Education Plan
INDI: International Neuroimaging Data-sharing Initiative
IQ: intelligence quotient
IRB: Institutional Review Board
KBIT: Kaufman Brief Intelligence Test
KSADS: Schedule for Affective Disorders and Schizophrenia for Children
LORIS: Longitudinal Online Research and Imaging System
MRI: magnetic resonance imaging
NIH: National Institutes of Health
NKI: Nathan Kline Institute
NY: New York
NYC: New York City
PHI: protected health information
PIQ: performance intelligence quotient
PPVT: Peabody Picture Vocabulary Test
QA: quality assurance
QAP: quality assurance plan
RUBIC: Rutgers University Brain Imaging Center
RV: research vehicle
SMI: SensoMotoric Instruments sMRI: structural magnetic resonance imaging
TOWRE: Test of Word Reading Efficiency
VIQ: verbal intelligence quotient
WAIS-IV: Wechsler Adult Intelligence Scale, 4th Edition
WASI: Wechsler Abbreviated Scale of Intelligence
WIAT-III: Wechsler Individual Achievement Test, 3rd Edition
WISC-V: Wechsler Intelligence Scale for Children, 5th Edition

### ETHICS APPROVAL AND CONSENT TO PARTICIPATE

All experimental procedures were performed with the approval of the Chesapeake Institutional Review Board and only after informed consent was obtained.

### CONSENT FOR PUBLICATION

All participants consented to have their data shared.

### COMPETING INTERESTS

The authors declare that they have no competing interests.

### FUNDING

The HBN (http://www.healthybrainnetwork.org) and its initiatives are supported by philanthropic contributions from the following individuals, foundations and organizations: Margaret Bilotti; Brooklyn Nets; Agapi and Bruce Burkard; James Chang; Phyllis Green and Randolph Cōwen; Grieve Family Fund; Susan Miller and Byron Grote; Sarah and Geoff Gund; George Hall; Jonathan M. Harris Family Foundation; Joseph P. Healey; The Hearst Foundations; Eve and Ross Jaffe; Howard & Irene Levine Family Foundation; Rachael and Marshall Levine; George and Nitzia Logothetis; Christine and Richard Mack; Julie Minskoff; Valerie Mnuchin; Morgan Stanley Foundation; Amy and John Phelan; Roberts Family Foundation; Jim and Linda Robinson Foundation, Inc.; The Schaps Family; Zibby Schwarzman; Abigail Pogrebin and David Shapiro; Stavros Niarchos Foundation; Preethi Krishna and Ram Sundaram; Amy and John Weinberg; Donors to the 2013 Child Advocacy Award Dinner Auction; Donors to the 2012 Brant Art Auction

## AUTHOR CONTRIBUTIONS

### Conception and Experimental Design

CA, LMA, JE, MPM, RCC, HSK, NL, CL, AM, KRM, JM, AN, KRP, FXC, AK, TP, BLL, SC, SPK, LCP, BK, JG, MH, GM

### Implementation and Logistics

AA, CA, LMA, JE, MPM, AM, NVP, KF, SL, BOH, MB, NGV, KK*l*, YO, BS, RT, AK, TP, BLL, JC

### Data Collection

CA, KF, AA, MK, SL, BOH, JA, BB, AB, HB, VC, NC, EC, DC, ME, BF, NGV, GG, CG, EH, SH, DK, KK, KK*l*, EK, DK, GM, RN, AN, YO, AR, AR*e*, TS, BS, RW, AW, AY

### Data Informatics

LMA, MPM, RCC, LA, BK

### Data Analysis

LMA, LA, MPM

### Drafting of the Manuscript

LMA, MPM

### Critical Review and Editing of the Manuscript

All authors contributed to the critical review and editing of the manuscript.

## ACKNOWLEDGEMENTS

We thank the Communications, Development, Finance, and Human Resource teams at the Child Mind Institute (past and present) for their endless support, and the CMI Executive Team - particularly, Elizabeth Planet, Natalie Cumberbatch, Brett Datkin, Dwayne Flinchum, and David Rivera; also the Child Mind Institute Scientific Research Council (https://childmind.org/our-research/scientific-research-council/) for their guidance and critical feedback in the planning of the Healthy Brain Network. Tammy Vanderwaal and Uri Hasson for their consultation in the selection of movies for the natural viewing paradigms; Simon Kelly for his assistance with devising the EEG battery; Megan Horton for advising us to add the collection of baby teeth; Antonio Convit for information regarding assessments of body composition; Stan Colcombe for information regarding fitness assessments; and Michael Michaelides for advising us to add hair samples for metals. Additionally, we would like to thank Joan Kaufman and Ken Kobak for providing access to the newly developed computerized KSADS and Ted Satterthwaite for helpful comments on the manuscript during its preparation.

We also acknowledge and thank Staten Island Borough President James Oddo, Staten Island Health and Wellness Director Dr. Ginny Mantello, and New York State Senator Andrew J. Lanza and his team (specifically William Matarazzo and Anthony Reinhart) for their guidance in developing strong partnerships throughout Staten Island, and their continued support of the project. We would also like to express our sincere gratitude to the mental health organizations, service providers, and clinicians across Staten Island, and NYC at large, who continue to work with our staff and refer participants to the project.

## AUTHOR DETAILS

1. Center for the Developing Brain, Child Mind Institute, New York, NY, USA

2. Center for Biomedical Imaging and Neuromodulation, Nathan S. Kline Institute for Psychiatric Research, New York, NY, USA

3. Center for Autism and the Developing Brain, Weill Cornell Medicine, New York-Presbyterian, White Plains, NY, USA

4. Columbia University Medical Center, New York, NY, USA

5. Division of Child and Adolescent Psychiatric Research, Nathan Kline Institute for Psychiatric Research, Orangeburg, NY, USA

6. Genetic Epidemiology Research Branch, Intramural Research Program, National Institute of Mental Health, Bethesda, MD, USA

7. Mercy College, Dobbs Ferry, NY, USA

8. New York University Langone Medical Center, New York, NY, USA

9. Rotman Research Institute, Baycrest, Toronto, Canada

10. The Child Study Center at NYU Langone Medical Center, New York, NY, USA

11. University of California San Francisco, San Francisco, CA, USA

12. Departments of Psychology and Psychiatry, University of Toronto, Toronto, Canada

13. Universität Zürich, Zürich, Switzerland

14. Yale University, New Haven, CT, USA

15. City University of New York, NY, USA

16. Office of Staten Island Borough President, Staten Island, NY, USA

17. Icahn School of Medicine at Mount Sinai, New York, NY, USA

18. New York State Institute for Basic Research in Developmental Disabilities, Staten Island, NY, USA

19. School of Electrical and Electronic Engineering, University College Dublin, Dublin, Ireland

## REFERENCES

1. Kessler, R. C. et al. Lifetime prevalence and age-of-onset distributions of DSM-IV disorders in the National Comorbidity Survey Replication. Arch. Gen. Psychiatry 62, 593–602 (2005).

2. Di Martino, A. et al. Unraveling the miswired connectome: a developmental perspective. Neuron 83, 1335–1353 (2014).

3. Insel, T. R. & Cuthbert, B. N. Medicine. Brain disorders? Precisely. Science 348, 499–500 (2015).

4. Kapur, S., Phillips, A. G. & Insel, T. R. Why has it taken so long for biological psychiatry to develop clinical tests and what to do about it? Mol. Psychiatry 17, 1174–1179 (2012).

5. Milham, M. P., Craddock, R. C. & Klein, A. Clinically useful brain imaging for neuropsychiatry: How can we get there? Depress. Anxiety (2017). doi: 10.1002/da.22627

6. Cuthbert, B. N. & Insel, T. R. Toward the future of psychiatric diagnosis: the seven pillars of RDoC. BMC Med. 11, (2013).

7. Insel, T. et al. Research domain criteria (RDoC): toward a new classification framework for research on mental disorders. Am. J. Psychiatry 167, 748–751 (2010).

8. Fair, D. A., Bathula, D., Nikolas, M. A. & Nigg, J. T. Distinct neuropsychological subgroups in typically developing youth inform heterogeneity in children with ADHD. Proc. Natl. Acad. Sci. U. S. A. 109, 6769–6774 (2012).

9. Miranda-Dominguez, O. et al. Connectotyping: model based fingerprinting of the functional connectome. PLoS One 9, e111048 (2014).

10. Van Dam, N. T. et al. Data-Driven Phenotypic Categorization for Neurobiological Analyses: Beyond DSM-5 Labels. Biol. Psychiatry (2016). doi: 10.1016/j.biopsych.2016.06.027

11. Paus, T. Population neuroscience: Why and how. Hum. Brain Mapp. 31, 891–903 (2010).

12. Nooner, K. B. et al. The NKI-Rockland Sample: A Model for Accelerating the Pace of Discovery Science in Psychiatry. Front. Neurosci. 6, 152 (2012).

13. Kaufman, J. et al. Schedule for Affective Disorders and Schizophrenia for School-Age Children-Present and Lifetime Version (K-SADS-PL): initial reliability and validity data. J. Am. Acad. Child Adolesc. Psychiatry 36, 980–988 (1997).

14. Lord, C. et al. Autism diagnostic observation schedule‐‐2nd edition (ADOS-2). Los Angeles, CA: Western Psychological Corporation (2012).

15. Semel, E. M., Wiig, E. H. & Secord, W. *CELF3: clinical evaluation of language fundamentals*. (Psychological Corporation, Harcourt Brace, 1995).

16. Wechsler, D. Wechsler intelligence scale for children‐‐Fourth Edition (WISC-IV). San Antonio, TX: The Psychological Corporation (2003).

17. Kaufman, A. S. & Kaufman, N. L. in *Encyclopedia of Special Education* (John Wiley & Sons, Inc., 2013). doi: 10.1002/9781118660584.ese1325

18. Wechsler, D. Wechsler Adult Intelligence Scale‐‐Fourth edition (WAIS-IV), Australian and New Zealand Language Adaptation. San Antonio, TX: NCS Pearson Inc (2008).

19. Test, W. D. W. I. A. (WIAT II). The Psychological Corp, London (2005).

20. Goldman, R. GFTA-3: Goldman-Fristoe test of articulation 3 [assessment instrument]. (2015).

21. Wagner, R. K., Torgesen, J. K., Rashotte, C. A. & Pearson, N. A. Comprehensive Test of Phonological Processing: CTOPP2. (2013).

22. Torgesen, J. K., Wagner, R. K. & Rashotte, C. A. *TOWRE: Test of word reading efficiency*. (Psychological Corporation, 1999).

23. Wiig, E. H., Semel, E. M. & Secord, W. *CELF 5: Clinical Evaluation of Language Fundamentals*. (Pearson/PsychCorp, 2003).

24. Williams, K. T. *EVT-2: Expressive vocabulary test*. (Pearson Assessments, 2007).

25. Dunn, L. M. & Dunn, D. M. *PPVT-4: Peabody picture vocabulary test*. (Pearson Assessments, 2007).

26. Achenbach, T. M. Integrative manual for the 1991 CBCL/4-18, YSR, and TRF profiles. Burlington, VT: University of Vermont, Department of Psychiatry (1991).

27. Gershon, R. C. et al. NIH toolbox for assessment of neurological and behavioral function. Neurology 80, S2–6 (2013).

28. Koffarnus, M. N. & Bickel, W. K. A 5-trial adjusting delay discounting task: accurate discount rates in less than one minute. Exp. Clin. Psychopharmacol. 22, 222–228 (2014).

29. The Cooper Institute. *Fitnessgram and Activitygram Test Administration Manual-Updated* 4th Edition. (Human Kinetics, 2010).

30. Langer, N. et al. A resource for assessing information processing in the developing brain using EEG and eye tracking. Sci Data 4, 170040 (2017).

31. Moeller, S. et al. Multiband multislice GE-EPI at 7 tesla, with 16-fold acceleration using partial parallel imaging with application to high spatial and temporal whole-brain fMRI. Magn. Reson. Med. 63, 1144–1153 (2010).

32. Deoni, S. C. L., Dean, D. C., 3rd, O’Muircheartaigh, J., Dirks, H. & Jerskey, B. A. Investigating white matter development in infancy and early childhood using myelin water faction and relaxation time mapping. Neuroimage 63, 1038–1053 (2012).

33. Sathian, K. et al. Dual pathways for haptic and visual perception of spatial and texture information. Neuroimage 57, 462–475 (2011).

34. LaConte S, Glielmi C, Heberlein K, Hu X. Verifying visual fixation to improve fMRI with predictive eye estimation regression (PEER). Proc. Intl. Soc. Magn. Reson. Med. 3438 (2007).

35. LaConte, S., Peltier, S., Heberlein, K. & Hu, X. Predictive eye estimation regression (PEER) for simultaneous eye tracking and fMRI. in Proc. Intl. Soc. Magn. Reson. Med 2808, (2006).

36. Sano, A. et al. Recognizing Academic Performance, Sleep Quality, Stress Level, and Mental Health using Personality Traits, Wearable Sensors and Mobile Phones. Int Conf Wearable Implant Body Sens Netw 2015, (2015).

37. Cohen, A. S., Renshaw, T. L., Mitchell, K. R. & Kim, Y. A psychometric investigation of ‘macroscopic’ speech measures for clinical and psychological science. Behav Res 48, 475–486 (2016).

38. Ghosh, S. S., Ciccarelli, G., Quatieri, T. F. & Klein, A. Speaking one’s mind: Vocal biomarkers of depression and Parkinson disease. J. Acoust. Soc. Am. 139, 2193–2193 (2016).

39. Markova, D. et al. Age- and sex-related variations in vocal-tract morphology and voice acoustics during adolescence. Horm. Behav. 81, 84–96 (2016).

40. Rowlands, A. V., Yates, T., Davies, M., Khunti, K. & Edwardson, C. L. Raw Accelerometer Data Analysis with GGIR R-package: Does Accelerometer Brand Matter? Med. Sci. Sports Exerc. 48, 1935–1941 (2016).

41. Berkovitz, B. K. B., Holland, G. R. & Moxham, B. J. *Oral anatomy, histology and embryology*. 398 (Elsevier, 2016).

42. Arora, M. et al. Spatial distribution of lead in human primary teeth as a biomarker of pre- and neonatal lead exposure. Sci. Total Environ. 371, 55–62 (2006).

43. Arora, M. et al. Determining prenatal, early childhood and cumulative long-term lead exposure using micro-spatial deciduous dentine levels. PLoS One 9, e97805 (2014).

44. Arora, M. et al. Determining fetal manganese exposure from mantle dentine of deciduous teeth. Environ. Sci. Technol. 46, 5118–5125 (2012).

45. Andra, S. S., Austin, C. & Arora, M. The tooth exposome in children’s health research. Curr. Opin. Pediatr. 28, 221–227 (2016).

46. Austin, C. et al. Uncovering system-specific stress signatures in primate teeth with multimodal imaging. Sci. Rep. 6, 18802 (2016).

47. Wechsler, D. Manual for the Wechsler abbreviated intelligence scale (WASI). San Antonio, TX: The Psychological Corporation (1999).

48. McCrimmon, A. W. & Climie, E. A. Wechsler D., Wechsler Individual Achievement Test—Third Edition. San Antonio, TX: NCS Pearson, 2009. Canadian Journal of School Psychology 26, 148–156 (2011).

49. Andra, S. S., Austin, C., Wright, R. O. & Arora, M. Reconstructing pre-natal and early childhood exposure to multi-class organic chemicals using teeth: Towards a retrospective temporal exposome. Environ. Int. 83, 137–145 (2015).

50. Simony, E. et al. Dynamic reconfiguration of the default mode network during narrative comprehension. Nat. Commun. 7, 12141 (2016).

51. Guntupalli, J. S. et al. A Model of Representational Spaces in Human Cortex. Cereb. Cortex 26, 2919–2934 (2016).

52. Birmaher, B. et al. Psychometric properties of the Screen for Child Anxiety Related Emotional Disorders (SCARED): a replication study. J. Am. Acad. Child Adolesc. Psychiatry 38, 1230–1236 (1999).

53. Mennes, M., Biswal, B. B., Castellanos, F. X. & Milham, M. P. Making data sharing work: the FCP/INDI experience. Neuroimage 82, 683–691 (2013).

54. Gorgolewski, K. J. et al. The brain imaging data structure, a format for organizing and describing outputs of neuroimaging experiments. Sci Data 3, 160044 (2016).

55. Barratt, W. The Barratt simplified measure of social status (BSMSS). Terre Haute, IN: Indiana State University (2006).

56. Zuo, X.-N. et al. An open science resource for establishing reliability and reproducibility in functional connectomics. Sci Data 1, 140049 (2014).

57. Di Martino, A. et al. Enhancing studies of the connectome in autism using the autism brain imaging data exchange II. Sci Data 4, 170010 (2017).

58. Frontiers | The Preprocessed Connectomes Project Quality Assessment Protocol - a resource for measuring the quality of MRI data. Available at: http://www.frontiersin.org/10.3389/conf.fnins.2015.91.00047/event_abstract. (Accessed: 11th June 2017)

59. Jenkinson, M., Bannister, P., Brady, M. & Smith, S. Improved optimization for the robust and accurate linear registration and motion correction of brain images. Neuroimage 17, 825–841 (2002).

60. Power, J. D., Schlaggar, B. L. & Petersen, S. E. Recent progress and outstanding issues in motion correction in resting state fMRI. Neuroimage 105, 536–551 (2015).

61. Ciric, R. et al. Benchmarking of participant-level confound regression strategies for the control of motion artifact in studies of functional connectivity. Neuroimage (2017). doi: 10.1016/j.neuroimage.2017.03.020

62. Power, J. D., Barnes, K. A., Snyder, A. Z., Schlaggar, B. L. & Petersen, S. E. Spurious but systematic correlations in functional connectivity MRI networks arise from subject motion. Neuroimage 59, 2142–2154 (2012).

63. Vanderwal, T., Kelly, C., Eilbott, J., Mayes, L. C. & Castellanos, F. X. Inscapes: A movie paradigm to improve compliance in functional magnetic resonance imaging. Neuroimage 122, 222–232 (2015).

64. Delorme, A. & Makeig, S. EEGLAB: an open source toolbox for analysis of single-trial EEG dynamics including independent component analysis. J. Neurosci. Methods 134, 9–21 (2004).

65. Gershon, J. A meta-analytic review of gender differences in ADHD. J. Atten. Disord. 5, 143–154 (2002).

66. Holmes, A. J. et al. Brain Genomics Superstruct Project initial data release with structural, functional, and behavioral measures. Sci Data 2, 150031 (2015).

67. Satterthwaite, T. D. et al. An improved framework for confound regression and filtering for control of motion artifact in the preprocessing of resting-state functional connectivity data. Neuroimage 64, 240–256 (2013).

68. Pruim, R. H. R. et al. ICA-AROMA: A robust ICA-based strategy for removing motion artifacts from fMRI data. Neuroimage 112, 267–277 (2015).

69. Behzadi, Y., Restom, K., Liau, J. & Liu, T. T. A component based noise correction method (CompCor) for BOLD and perfusion based fMRI. Neuroimage 37, 90–101 (2007).

70. Chai, X. J., Castañón, A. N., Ongür, D. & Whitfield-Gabrieli, S. Anticorrelations in resting state networks without global signal regression. Neuroimage 59, 1420–1428 (2012).

71. Yan, C.-G., Craddock, R. C., Zuo, X.-N., Zang, Y.-F. & Milham, M. P. Standardizing the intrinsic brain: towards robust measurement of inter-individual variation in 1000 functional connectomes. Neuroimage 80, 246–262 (2013).

72. Betzel, R. F. et al. Generative models of the human connectome. Neuroimage 124, 1054–1064 (2016).

73. Bezgin, G., Solodkin, A., Bakker, R., Ritter, P. & McIntosh, A. R. Mapping complementary features of cross-species structural connectivity to construct realistic ‘Virtual Brains’. Hum. Brain Mapp. 38, 2080–2093 (2017).

74. Klein, A. et al. Mindboggling morphometry of human brains. PLoS Comput. Biol. 13, e1005350 (2017).

75. Vanderwal, T. et al. Individual differences in functional connectivity during naturalistic viewing conditions. (2016). doi: 10.1101/084665

76. Kauttonen, J., Hlushchuk, Y. & Tikka, P. Optimizing methods for linking cinematic features to fMRI data. Neuroimage 110, 136–148 (2015).

77. Dmochowski, J. P. et al. Audience preferences are predicted by temporal reliability of neural processing. Nat. Commun. 5, (2014).

78. Dubois, J. & Adolphs, R. Building a Science of Individual Differences from fMRI. Trends Cogn. Sci. 20, 425–443 (2016).

79. Wang, J. et al. Test-retest reliability of functional connectivity networks during naturalistic fMRI paradigms. Hum. Brain Mapp. 38, 2226–2241 (2017).

80. O’Connor, D. et al. The Healthy Brain Network Serial Scanning Initiative: a resource for evaluating inter-individual differences and their reliabilities across scan conditions and sessions. Gigascience 6, 1–14 (2017).

81. Poldrack, R. A. et al. Long-term neural and physiological phenotyping of a single human. Nat. Commun. 6, 8885 (2015).

82. Fayers, P. Item Response Theory for Psychologists. Qual. Life Res. 13, 715–716 (2004).

83. Embretson, S. E. & Reise, S. P. *Item Response Theory*. (Psychology Press, 2013).

84. Kim-O, M.-A. & Embretson, S. E. in *Handbook of Behavioral Medicine* 113–123 (2010). doi: 10.1007/978-0-387-09488-5_9

85. Magnotta, V. A., Friedman, L. & FIRST BIRN. Measurement of Signal-to-Noise and Contrast-to-Noise in the fBIRN Multicenter Imaging Study. J. Digit. Imaging 19, 140–147 (2006).

86. Mortamet, B. et al. Automatic quality assessment in structural brain magnetic resonance imaging. Magn. Reson. Med. 62, 365–372 (2009).

87. Atkinson, D., Hill, D. L., Stoyle, P. N., Summers, P. E. & Keevil, S. F. Automatic correction of motion artifacts in magnetic resonance images using an entropy focus criterion. IEEE Trans. Med. Imaging 16, 903–910 (1997).

88. Friedman, L., Glover, G. H., Krenz, D., Magnotta, V. & FIRST BIRN. Reducing inter-scanner variability of activation in a multicenter fMRI study: role of smoothness equalization. Neuroimage 32, 1656–1668 (2006).

89. Giannelli, M., Diciotti, S., Tessa, C. & Mascalchi, M. Characterization of Nyquist ghost in EPI-fMRI acquisition sequences implemented on two clinical 1.5 T MR scanner systems: effect of readout bandwidth and echo spacing. J. Appl. Clin. Med. Phys. 11, 3237 (2010).

90. Nichols, T. E. Standardizing DVARS. (2012). Available at: http://blogs.warwick.ac.uk/nichols/entry/standardizing_dvars.

91. Cox, R. W. AFNI: software for analysis and visualization of functional magnetic resonance neuroimages. Comput. Biomed. Res. 29, 162–173 (1996).

